# Adaptive ratchets and the evolution of molecular complexity

**DOI:** 10.1101/2021.11.18.469165

**Authors:** Tom Röschinger, Roberto Morán Tovar, Simone Pompei, Michael Lässig

## Abstract

Biological systems have evolved to amazingly complex states, yet we do not understand in general how evolution operates to generate increasing genetic and functional complexity. Molecular recognition sites are short genome segments or peptides binding a cognate recognition target of sufficient sequence similarity. Such sites are simple, ubiquitous modules of sequence information, cellular function, and evolution. Here we show that recognition sites, if coupled to a time-dependent target, can rapidly evolve to complex states with larger code length and smaller coding density than sites recognising a static target. The underlying fitness model contains selection for recognition, which depends on the sequence similarity between site and target, and a uniform cost per unit of code length. Site sequences are shown to evolve in a specific adaptive ratchet, which produces selection of different strength for code extensions and compressions. Ratchet evolution increases the adaptive width of evolved sites, accelerating the adaptation to moving targets and facilitating refinement and innovation of recognition functions. We apply these results to the recognition of fast-evolving antigens by the human immune system. Our analysis shows how molecular complexity can evolve as a collateral to selection for function in a dynamic environment.

## Introduction

Darwin’s principle states that evolution progresses by mutations and selection. This principle describes a dynamical pathway to organisms of high fitness, the efficacy of which is modulated by other, stochastic evolutionary forces. In microbial systems, for example, fitness can increase drastically even on time scales of laboratory evolution experiments; these processes are carried by multiple mutations and occur repeatably in parallel-evolving populations [1]. In a similar way, increasing complexity appears to be a ubiquitous feature of biological evolution. Already the onset of evolution requires the complexity of biopolymers to grow above a minimum required for autocatalytic replication [2, 3]. In macro-evolution, a prominent example is the increase of genome and regulatory complexity in the transition from prokaryotic to multicellular eukaryotic organisms [4]. Nevertheless, the evolutionary forces behind complexity have remained controversial [5]. From a classical Darwinian perspective, organisms evolve complex functions because they adapt to complex environments; that is, complexity increases by positive selection [6]. However, producing and maintaining complex features also requires a physiological machinery. This generates a fitness cost of complexity, the relative weight of which is often unknown. Recent genomic and functional experiments show that molecular complexity can arise by neutral evolution and be maintained by selection, even if there is no discernible fitness benefit [7–15]. On much shorter time scales, humans and other vertebrates evolve complex antibody repertoires during their lifetime; these antibodies encode a memory of past infections and serve in the immune defense against future infections by similar antigens [16, 17]. Is there a universal force that drives evolving systems to higher complexity, much as selection acts to increase fitness, or are the explanations for complex outcomes as complex as the subject matter? This question has no satisfactory answer to date. We lack a general definition of complexity and a unifying theory that predicts when complexity is expected to evolve.

In this paper, we study the evolution of complexity for recognition sites in DNA or proteins. By binding a cognate molecular target, such sites perform specific regulatory, enzymatic, or signaling tasks that feed into larger functional networks of the cell. Recognition sites are simple, ubiquitous, and often quantitatively understood units of molecular function, making them an ideal subject of our study. Most importantly, a single molecular phenotype, the binding affinity to the recognition target, characterizes the functionality of a site and is the target of natural selection. At the sequence level, recognition sites are approximately digital units of molecular information. The binding specificity of the target is often summarized in a sequence motif that lists the preferred nucleotides or amino acids at each position of the recognition site. Functional sites are biased towards sequences matching the target, which typically generates a relative information, or entropy loss, of order 1 bit per unit of functional sequence [18,19]. The code length of a recognition site can be taken as a measure of its molecular complexity. This measure is quite variable across different classes of recognition sites. Strong transcription factor binding sites in prokaryotes have typical lengths ~ 10 base pairs; these sites are compact functional units close to the minimal coding length required for function. This lower bound is comparable to the algorithmic complexity of a computer program predicting recognition of the target [20,21]. Weak binding sites, often acting together with adjacent sites, spread this information to larger code length and lower coding density. Prokaryotic RNA polymerase binding sites contain about 40 base pairs long, a subset of these positions carries a highly conserved recognition code [22,23]. More complex transcription units containing low-affinity sites are also common in eukaryotes [24]. Similarly, human immune recognition can be carried by compact T cell receptors binding to antigenic peptides of code length < 10 amino acids; more complex immune responses can involve multiple antibody lineages interacting with their cognate antigenic sites [25]. Immune recognition systems have recently been discussed as targets of host-pathogen co-evolution and eco-evolutionary control [26–28]. These examples contain the main question of this paper. When is the recognition encoded in sites close to minimal code length needed for specificity? Conversely, when do we find longer and fuzzier sites, where higher molecular complexity is associated with the same function?

Recognition sites and their complexity are the outcome of evolution. These dynamics has three kinds of elementary steps. Point mutations of the recognition site update individual letters and may affect the coding density; code extensions and compressions also change the length of the site. Moreover, in most instances, the target of recognition evolves itself. Target changes can be independent of the recognition function, as it is often the case for proteins with multiple functions. They can also be directed to change recognition; for example, the evolution of transcription factors can update regulatory networks [29–33]. On shorter time scales, a prominent example is escape mutations of antigens from immune recognition by their hosts. Natural selection acts on the recognition function, which depends on the target binding affinity Δ*G*, but is blind to details of site and target sequence. This has an important consequence: at a given level of Δ*G*, the molecular complexity of sites does not carry any direct fitness benefit. Moving targets generate time-dependent selection and, if the rates of change are mutually in tune, induce a continual adaptive evolution of their cognate sites. Coevolving sites can maintain recognition, albeit at a lower affinity Δ*G* than for static targets, because the adaptation of the site sequence always lags behind the evolution of the target [34]. Together, these dynamics shape sequence architecture, information content, and functionality of recognition sites.

Here, we describe the evolution of recognition sites in a minimal, analytically solvable, yet bio-physically realistic model of function and fitness. The model maps two evolutionary pathways towards molecular complexity: sites become long and fuzzy when the cost of complexity is low or when the target of recognition is moving. In both cases, recognition sites evolve in a ratchet mode where opposite steps of code extension and compression do not have opposite selection coefficients. In the low-cost regime, the evolutionary ratchet operates by constraint: site extensions are near-neutral, site compressions are slowed down by selection on recognition. Comparable mechanisms have previously been discussed for genome and protein evolution [9,11,13–15]. In the moving-target regime, the ratchet becomes adaptive: all extension and compression steps are driven by positive selection on recognition. Adaptive ratchets generate a faster and more robust evolution of molecular complexity, which can overcome a substantial cost of complexity. We show that evolved sites have an increased susceptibility to selection, which facilitates adaptation to moving targets, the refinement of existing functions, and the adoption of new functions. As an application, we discuss the benefits of complexity in the immune response to fast-evolving antigens.

## Results

### Model for function and fitness

Recognition sites work in a complex molecular environment, which imposes stringent requirements for their functionality. First, spurious recognition generated by accidental binding of target molecules must be sufficiently rare. This sets a lower bound on the information content of recognition motifs, which can be defined as the Kulback-Leibler distance between the ensemble of target-bound sites and random sequence, *H*(*Q*|*P*_0_); see Methods. Second, functional sites must be target-bound at physiological concentrations of the target molecule, which sets a lower bound on the free energy gap to random sequence, Δ*G*. For transcriptional regulation in prokaryotes, these functionality requirements have been shown to broadly reproduce the observed architecture of minimally complex recognition sites [35–38]. They also inform the minimal model used in this study [39]: sites have a code length *ℓ* ≳ 10 and a sequence alphabet of *q* equiprobable letters (with *q* = 4 for nucleotides and *q* = 20 for amino acids); each sequence position has one matching letter and (*q* – 1) mismatches with a reduced free energy difference ΔΔ*G*/*k_B_T* = *ϵ*_0_ ~ 1. In this model, we can characterise recognition sites by just two summary variables: the code length *ℓ* and the number of matches, k, or equivalently, the coding density *γ* = *k*/*ℓ*.

Under conditions of thermodynamic equilibrium, recognition of the target depends on the reduced free energy gap Δ*G*/*k_B_T* = (*γ* – *γ*_0_)*ℓϵ*_0_, where *γ*_0_ = 1/*q* is the density of matches in random sequence. The recognition probability *R* is then a standard Hill function, *R*(*γ, ℓ*) = H(Δ*G* – Δ*G*_50_), where we assume a free energy Δ*G*_50_ ~ 10 *k_B_T* is required for half-binding under physiological conditions (Methods). In a biophysical fitness model, the recognition-dependent fitness of a site is taken to be proportional to the recognition probability, *F_r_*(*γ, ℓ*) = *f*_0_*R*(*γ, ℓ*). This type of fitness model generates specific fitness nonlinearities (epistasis) that can be traced in the evolutionary statistics of recognition sites [40, 41]. It has been broadly used for regulatory binding sites, protein interaction sites, and protein folding [12,36,37,42–46].

The landscape *F*(*γ, ℓ*) determines the selection acting on changes of the recognition site sequence. In the minimal model, a mutation from *k* to (*k* + 1) matches has the selection coefficient *s_γ_* = *F_r_*(*γ* + 1/ℓ,ℓ) – *F_r_*(*γ,ℓ*). As will become explicit in the following, most functional recognition sites are found on the upper branch of the Hill function just above the half-binding point (Fig. 1). In this region, we can approximate the Hill function by an exponential function, which shows that selection is proportional to the recognition error, *s_γ_* = *ϵ*_0_*f*_0_ Δ*R* with Δ*R* = 1 – *R*.

**Fig. 1:**
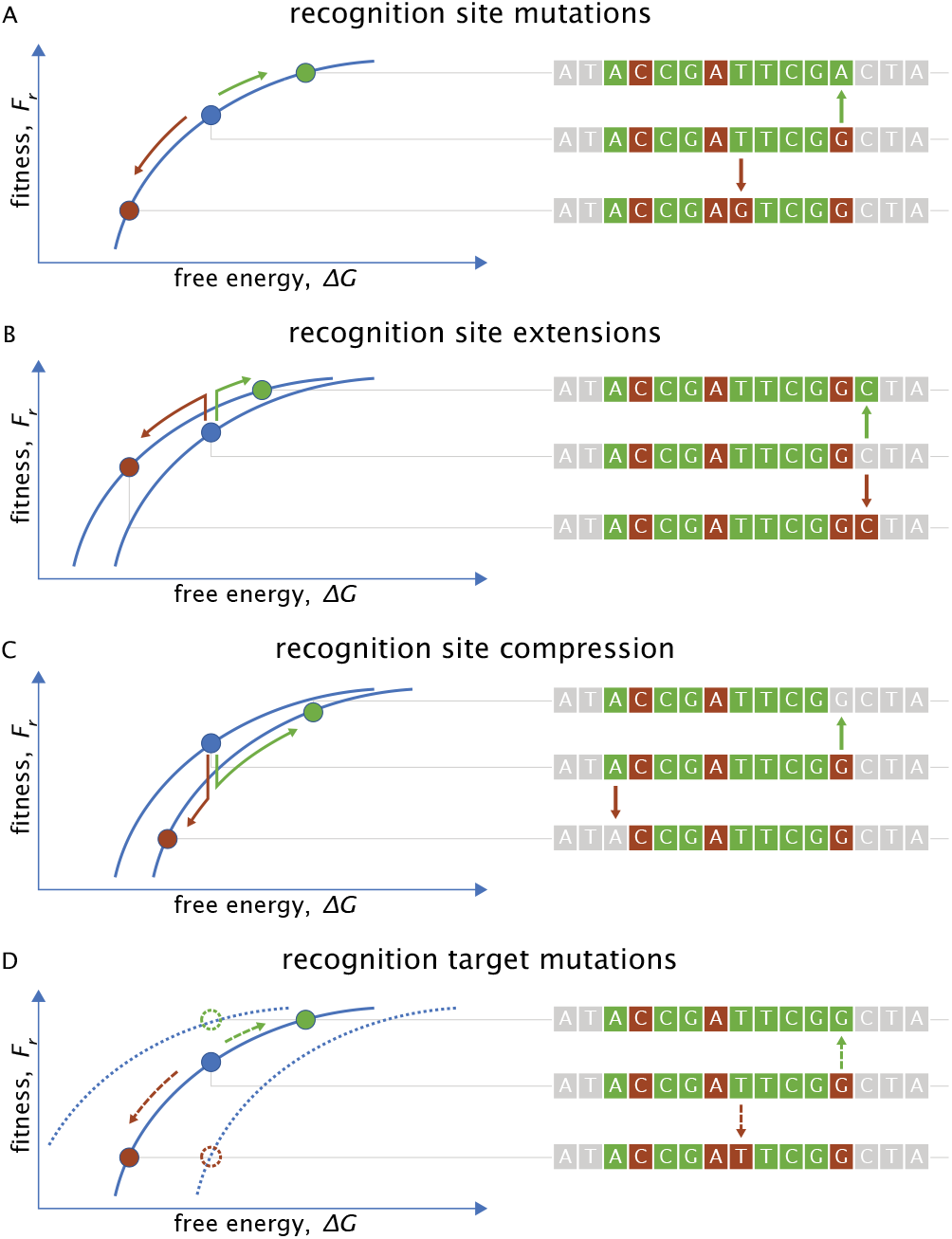
Evolution of molecular recognition. Evolutionary steps change the recognition code (green squares: target matches, purple squares: mismatches). These changes affect the free energy of recognition, Δ*G*, and the recognition fitness, *F_r_* (green arrows: beneficial changes, purple arrows: deleterious changes). (A) Point mutations of a recognition site add or remove one letter matching the target, *k* → *k* ± 1, at constant code length í. (B) Site extensions and (C) compressions add or remove one unit of length, *ℓ* → *ℓ* ± 1. (D) Recognition target mutations make the fitness of a given recognition site explicitly time-dependent (dotted lines) and act as effective site mutations with neutral rates (dashed arrows).

In a biophysical fitness model, selection acts on the recognition code only through the intermediate phenotype Δ*G*. This is an important prerequisite for the evolution of complexity: functionality of recognition imposes a lower bound on the code length, *ℓ* ≳ *ℓ*_0_ ≡ Δ*G*_50_/(*ϵ*_0_*k_B_T*), but does not generate an upper bound. Other components of cell physiology, including genome replication and competing molecular functions, are expected to constrain the code length of recognition sites. Here we describe this constraint by a linear fitness cost of genomic real estate, *F_c_* = −*c*_0_*ℓ*. Together, recognition fitness and cost of complexity define our minimal fitness model,

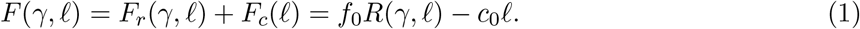

#### Evolutionary dynamics

We study the evolution of recognition sites in the low-mutation regime, where site sequences are updated sequentially by substitutions in an evolving population. The rates of these substitutions depend on the corresponding mutation rates and on selection coefficients scaled by an effective population size *N*, as given by Haldane’s formula [47] (Methods). Our minimal model contains three types of updating steps. First, point mutations of the site sequence occur at a homogeneous rate *μ* per unit of length (Fig. 1A). Hence, a recognition site with k matches and ℓ – *k* mismatches has a total rate *μ*_+_(*γ, ℓ*) = *μℓγγ*_0_/(1 – *γ*_0_) of beneficial mutations and a total rate *μ*___ (*γ, ℓ*) = *μℓ*(1 – *γ*) of deleterious mutations (Methods). These changes have selection coefficients ±*s_γ_*(*γ,ℓ*) and substitution rates *u*_±_(*γ,ℓ*), which depend on the fitness landscape *F*(*γ,ℓ*) introduced above.

Second, sites change by extension and compression steps, which add or remove one unit of sequence (Fig. 1BC). These changes, which affect the architecture of recognition, are assumed to occur at a much lower rate, *v* ≪ *μ*. Extensions add a random letter of background sequence to the site, whereas compressions remove a random letter of the existing site sequence. In the minimal model, beneficial and deleterious extensions occur with total rates *v*_++_(*γ,ℓ*) = *vγ*_0_ and *v*_+__(*γ,ℓ*) = *v*(1 – *γ*_0_), respectively; beneficial and deleterious compressions occur with total rates *v*_−+_(*γ,ℓ*) = *v*(1 – *γ*) and *v*_− −_ *γ, ℓ*) = *vγ*. These rates, together with the corresponding selection coefficients *s*_±±_(*γ,ℓ*) given by the fitness landscape *F*(*γ,ℓ*), determine the total substitution rates for extensions and compressions, *v*_+_(*γ,ℓ*) and *v*___(*γ,ℓ*) (Methods).

Third, the mutation target sequence changes at a rate *ρ* = *κμ* per unit of length (Fig. 1D). This rate is assumed to be comparable to the point mutation rate *μ* and to be generated by external factors independent of the recognition function. The selective effects of target changes take the form of a fitness seascape, generating an explicitly time-dependent fitness of a given recognition site sequence [48]. On the recognition-fitness map *F_r_*(*γ,ℓ*) and in the low mutation regime, target changes act as additional effective mutations that change the free energy Δ*G* and the recognition *R* with respect to the moving target. Experiments have shown that mutations in the DNA binding domain of LacI in *E. coli* can be reduced to a change in binding energy, similar to mutations in the DNA itself [49]. Importantly, the selective effects of target mutations are again given by the fitness landscape *F*(*γ,ℓ*) but their rates are not: beneficial and deleterious changes occur at neutral rates *ρ*_+_(*γ,ℓ*) = *ρℓγγ*_0_/(1 – *γ*_0_) and *ρ*___(*γ,ℓ*) = *ρℓ*(1 – *γ*), respectively, generating a net degradation of recognition and driving the adaptive evolution of recognition sites.

#### Stationary distributions

In an evolutionary steady state, recognition sites keep catching up with a moving target and maintain time-independent averages of binding affinity and recognition probability. In general, non-equilibrium processes in a nonlinear fitness landscape are quite complicated. Remarkably, however, the steady state of the minimal recognition model can be computed analytically for arbitrary target evolution rates *ρ* = *κμ*. We write the steady-state distribution of coding density and code length in an ensemble of parallel-evolving populations as the exponential of an evolutionary potential, *Q*(*γ,ℓ*) = exp[Φ(*γ,ℓ*)]. Because recognition site mutations occur at a much higher rate than extensions or compressions, the coding density relaxes to a stationary state between any two code length changes and the potential takes the form Φ(*γ,ℓ*) = Φ(*γ|ℓ*) + Φ(*ℓ*). The coding density component Φ(*γ|ℓ*) is built from ratios of substitution rates for beneficial code changes and their reverse deleterious changes at constant length, (*u*_+_(*γ′,ℓ*) + *ρ*_+_ (*γ*′,ℓ))/(*u*___(*γ*′ + 1/ℓ,ℓ) + *ρ*___(*γ′* + 1/ℓ,ℓ)). Similarly, the length component Φ(*ℓ*) can be written in terms of effective substitution rates for site extensions and their reverse compressions, 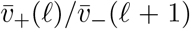; these rates are obtained by averaging the basic extension and compression rates over the conditional stationary distribution *Q*(*γ*|*ℓ*) or, in close approximation, by using the maximum-likelihood (ML) coding density *γ*\*(*ℓ*) (Methods, Fig. S1, Fig. S2). Furthermore, by weighting the distribution *Q*(*γ,ℓ*) with the recognition function *R*(*γ,ℓ*), we can define a distribution of functional codes, *Q_f_*(*γ,ℓ*). The full analytic solution is given in Methods, Eqs. [S16] - [S23].

Stationary distributions *Q_f_*(*γ,ℓ*) for different cost parameters and driving rates are shown in Fig. 2; global ML points (*γ*,ℓ*\*) are marked by dots. These plots display the central results of the paper. First, recognition sites for a static target (*κ* = 0) evolve increasing code length and decreasing coding density with decreasing cost of complexity, 2*N*_*c*_0__ (Fig. 2A); the cost-dependent ML values and widths of *γ* and *ℓ* are shown in Fig. 3AB. Second, recognition sites for a dynamic target evolve increasing molecular complexity with increasing driving rate, *κ* (Fig. 2C, Fig. 3DE). The analytical distributions *Q*(*γ,ℓ*) (Fig. 2AC) are in accordance with time averages over individual trajectories obtained by simulation (Fig. 2BD). As a function of the target mutation rate, we observe shifts in the substitution dynamics. For static targets (*κ* = 0), the stationary state shows a balance of beneficial and deleterious substitutions (Fig. 2B; Fig. 2D, left), indicating evolution at equilibrium. At strong driving (*κ* ≳ 1), substitutions are predominantly adaptations to target mutations (Fig. 2D, right). Hence, in this regime, enhanced code complexity is associated with deviations from evolutionary equilibrium.

**Fig. 2:**
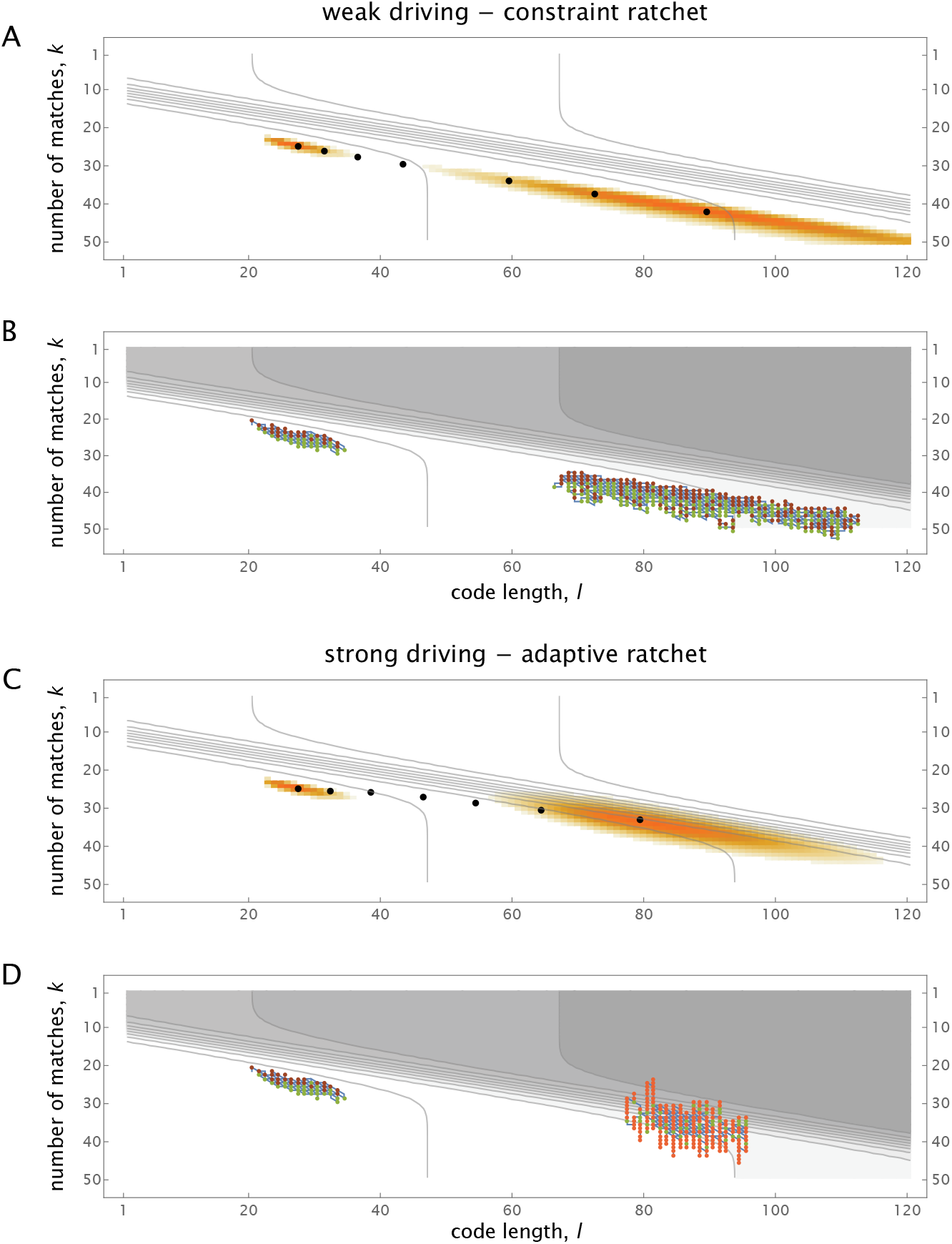
Statistics and evolutionary trajectories of recognition sites. (A, B) Joint stationary distributions of code length and number of matches for functional sites, *Q_f_*(*γ,ℓ*) with *γ* = *k/ℓ*, for different values of the scaled complexity cost, 2*N*_*c*_0__, and the driving rate, *κ*. Contours show the fitness landscape *F*(*γ,ℓ*), which is steepest at half-binding, *k* = *γ*_0_ℓ + *ℓ*_0_. (A) Weak-driving regime: distributions *Q_f_*(*γ,ℓ*) for 2*N*_*c*_0__ = 1 (left) and 2*N*_*c*_0__ = 0.1 (right) at *κ* = 0, maximum-likelihood points (*k*, ℓ*\*)(2*N*_*c*_0__, *κ* = 0) (black dots) for 2*N*_*c*_0__ = 1, 0.7,0.5, 0.35,0.2, 0.14, 0.1. (B) Crossover to the strong driving regime: distributions *Q_f_*(*γ,ℓ*) for *κ* = 0 (left) and *κ* = 45 (right) at 2*N*_*c*_0__ = 1, maximum-likelihood points (*k*,ℓ*\*)(2*N*_*c*_0__ = 1,*κ*) (black dots) for *κ* = 0, 0.2,0.5,1, 2, 5,10, 20,45. (C, D) Evolutionary paths, (*k,ℓ*)(*t*), with marked recognition site mutations (adaptive: green, deleterious: purple) and recognition target changes (red). Sample paths are shown (C) for 2*N*_*c*_0__ = 1 (left) and 2*N*_*c*_0__ = 0.1 (right) at *κ* = 0, and (D) for *κ* = 0 (left) and *κ* = 45 (right) at 2*N*_*c*_0__ = 1. Other evolutionary parameters: *γ*_0_ = 1/4, *ℓ*_0_ = 10, *ϵ*_0_ = 0.7, 2*Nf*_0_ = 400, *ν/μ* = 0.1; the evolutionary algorithm is detailed in Methods.

**Fig. 3:**
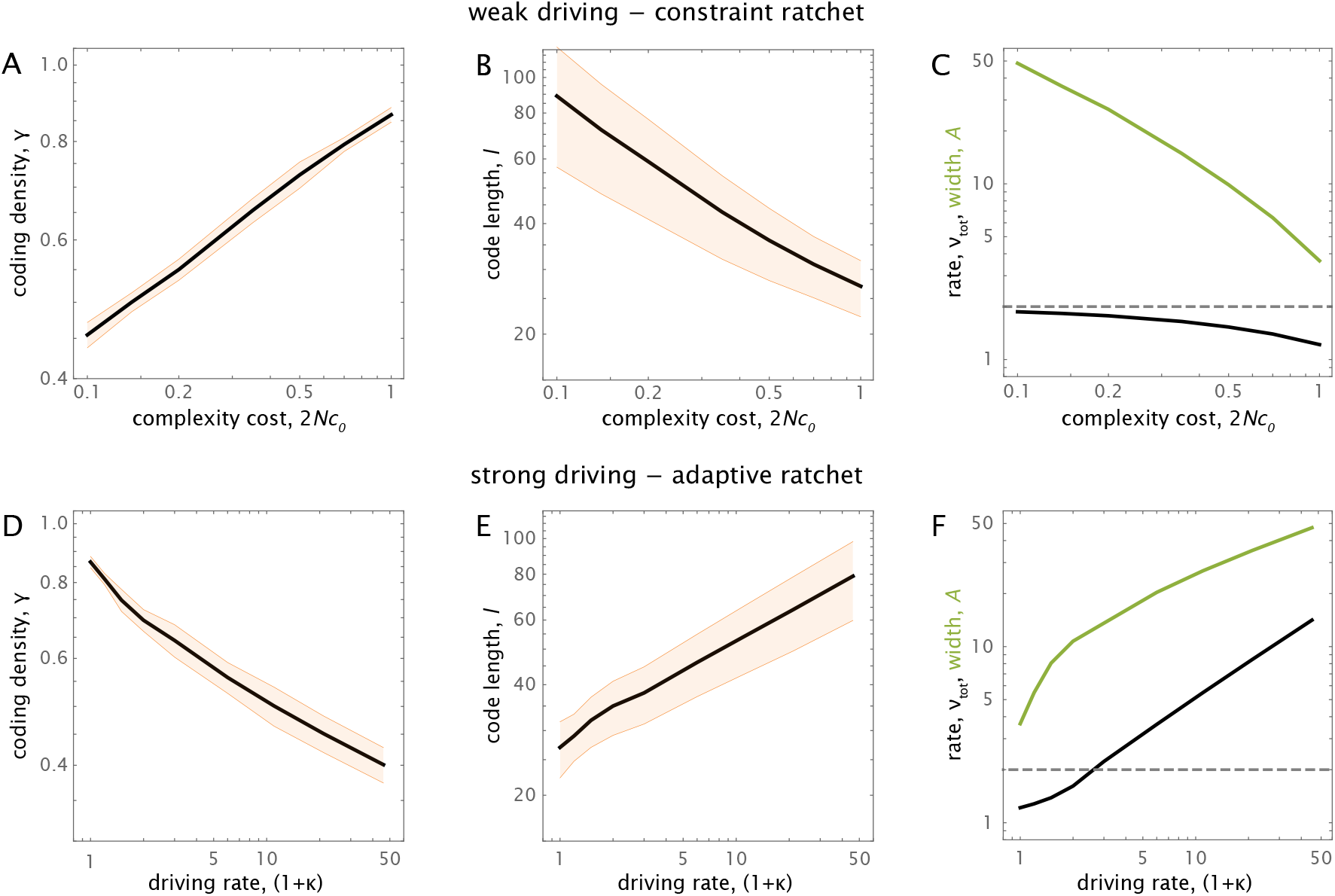
Complexity scaling. (A-C) Crossover to low cost in the weak-driving regime. (A) Coding density, ML value *γ*\*(*κ* = 0, *c*) (black), and standard deviation (orange). (B) Code length, ML value, *ℓ*\*(*κ* = 0, 2*N*_*c*_0__) (black), and standard deviation, Δ*ℓ*(*κ* = 0, 2*N*_*c*_0__) (orange). (C) ML adaptive width, *A*\*(*κ* = 0, 2*N*_*c*_0__) (green), and ML ratchet rate, 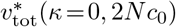 in units of *v* (black), compared to the neutral rate, 2*v* (dashed). (D-F) Crossover to strong-driving at unit cost. (D) Coding density, ML value *γ*\*(*κ*, 2*N*_*c*_0__ = 1) (black), and standard deviation (orange). (E) Code length, ML value, *ℓ*\*(*κ*, 2*N*_*c*_0__ = 1) (black), and standard deviation, Δ*ℓ*(*κ*, 2*N*_*c*_0__ = 1) (orange). (F) ML adaptive width, A*(κ, 2*N*_*c*_0__ = 1) (green), ML ratchet rate, 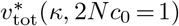 in units of *v* (black), and neutral rate, 2*v* (dashed). Evolutionary parameters as in Fig. 2.

#### Weak- and strong-driving regimes

The evolutionary potential Φ(*γ,ℓ*) simplifies at low and high target mutation rates, which allows for simple analytical estimates of the ML point (*γ*,ℓ\**) and gives insight in the evolutionary determinants of site complexity. We first recall that the code length *ℓ* and the conditional ML coding density, *γ*\*(*ℓ*), set the associated selection coefficient 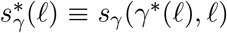, on the fitness landscape *F_r_*(*γ,ℓ*). At the global ML point (*γ*,ℓ\**), this relation can be written in the form

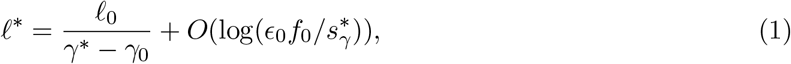

which says that the coding density settles on the upper branch of the recognition Hill function close to the inflection point.

The potential Φ(*γ*|*ℓ*) determines the ML coding density *γ*\*(*ℓ*). For weakly driven recognition sites (*κ* 1), we obtain the simple form Φ(*γ*|*ℓ*) = *S*(*γ,ℓ*) + 2*NF*(*γ,ℓ*)/(1 + *κ*) (Fig. S1), which is derived in Methods. Here, *N* is the effective population size and *S*(*γ,ℓ*) denotes the entropy of recognition sites, defined as the log number of sequence states of given coding density and code length. In the limit *κ* = 0, this potential reproduces the well-known Boltzmann-Gibbs form of a mutation-selection-drift equilibrium [36,50]. By maximizing Φ(*γ*|*κ,ℓ*), we obtain the weak-driving balance condition for coding density, 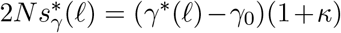 (Methods), which says that individual code letters evolve under marginal selection, 2*Ns*_γ_ ~ 1. In the equilibrium limit, this relation is accordance with general results for mutation-selection-drift-equilibria of quantitative traits [51–53]. With an appropriately rescaled effective population size, it holds even if clonal interference is the dominant noise in sequence evolution [54]. Marginal selection implies that the ML coding density can get close to but cannot reach 1, which explains the substantial sequence variation observed in families of transcription factor binding sites. In contrast, under strong driving (*κ* ≳ 1), maintaining recognition requires substantial selection on individual code letters. In this regime, deleterious mutations are gradually suppressed and the potential Φ(*γ*|*ℓ*) reflects the adaptive evolution of recognition sites in response to moving targets (Fig. 2D, Fig. S1). Again by maximizing Φ(*γ*|*ℓ*), we obtain the strong-driving balance relation for coding density, 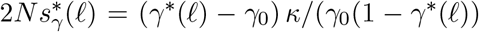 (Methods). Importantly, the weak- and strong-driving balance conditions are highly universal: the scaled selection coefficient 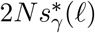 does not depend on the fitness amplitude *f_0_* or the effective population size *N*.

Given the conditional coding density *γ*\*(*ℓ*), the potential Φ(*ℓ*) determines the global ML point (*γ*,ℓ*\*) as a function of the evolutionary parameters. Maximizing this potential yields a balance condition for code length that relates selection on recognition, 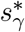, and the cost of complexity, *c*_0_. This condition has the weak- and strong-driving asymptotics 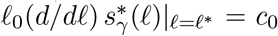 and 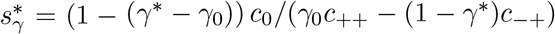, respectively, where *c*_++_ and *c*_−+_ are constants of order 0.1 – 1 (Methods). Together with the balance condition on coding density and Eq. [1], we obtain a closed expression for (*γ*,ℓ*\*). In the weak-driving regime, coding density and code length scale as powers of the cost, 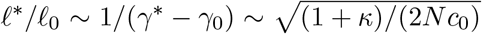. Hence, molecular complexity can be enhanced by a reduction in the long-term effective population size, for example, in the transition from prokaryotes to multicellular eukaryotes, as suggested previously for genome evolution [9]. For recognition sites, this pathway to complexity builds on the universality of selection for recognition, which causes the cost-benefit ratio 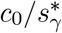 to decrease with N. A similar universality is found the strong-coupling regime: the ML point (*γ*,ℓ*\*) is the solution of a quadratic equation and depends only on the ratio 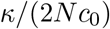 (Methods). In both regimes, the analytical closure is in accordance with the full solution (Fig. S3).

#### Ratchet evolution of code length

What kind of evolutionary mechanism generates the evolution of longer recognition codes? Consider first the long-term fitness effects of length changes: longer codes improve recognition but come with a cost of complexity. This tradeoff can be displayed by an effective fitness landscape for code length, 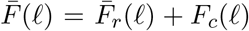, where the recognition component is defined by averaging *F*(*γ,ℓ*) over the conditional stationary distribution *Q*(*γ*|*ℓ*) (Fig. 4AC, Methods).

**Fig. 4:**
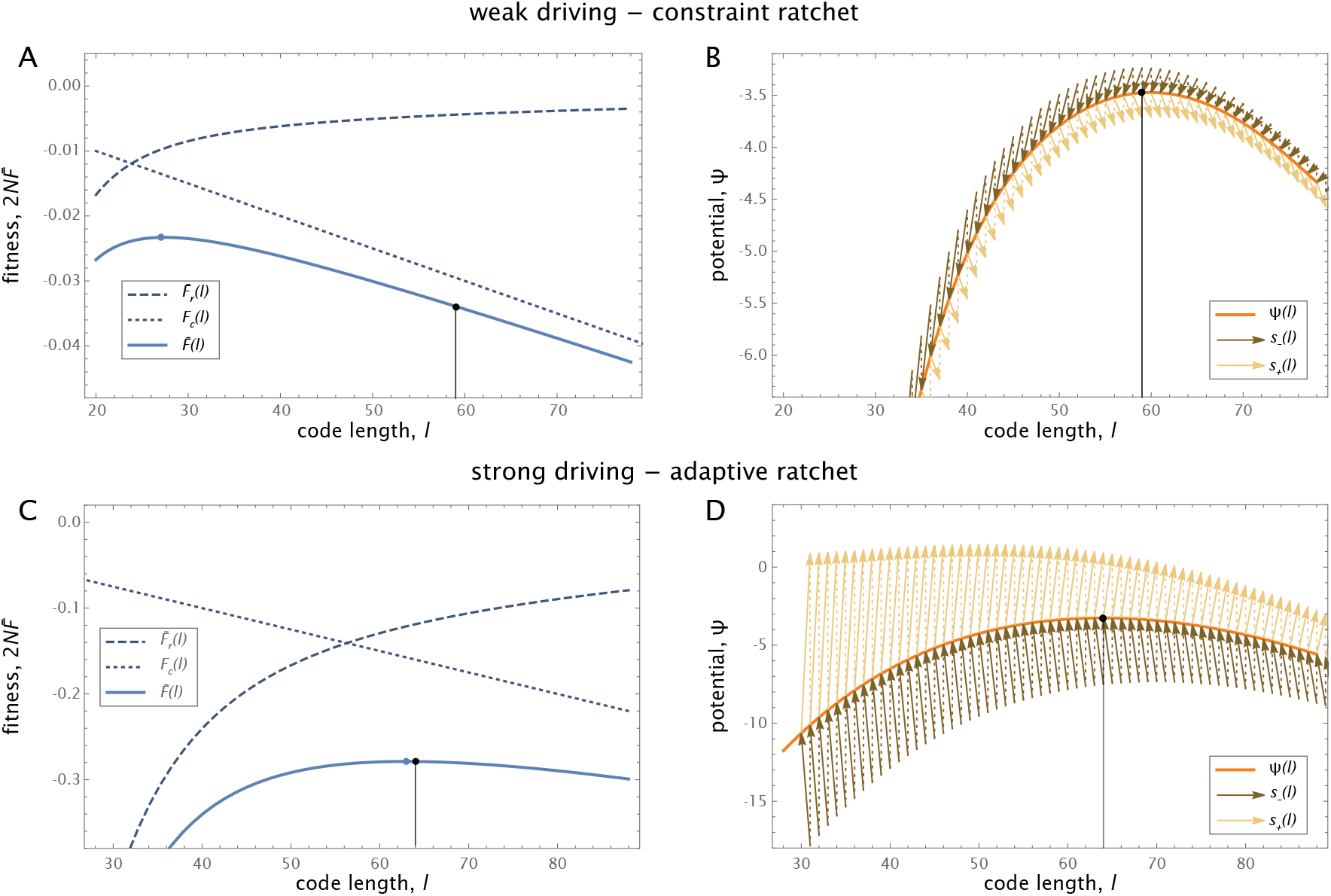
Ratchet evolution of code length. (A, C) Effective fitness landscape for code length, 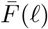 (blue solid), recognition component, 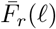 (blue dashed), and complexity cost, *F_c_*(*ℓ*) (blue dotted). ML values *γ*\* are marked by black dots; fitness values are relative to the maximum fitness and scaled by the effective population size *N*. (B, D) Evolutionary potential for length, Φ(*ℓ*) (orange). Arrows show ratchet selection coefficients for extension, *s*_+_(*ℓ*) (yellow), and for compression, *s*___(*ℓ*) (brown); see also Fig. S2. Dotted line segments guide the eye in joining adjacent steps. (A, B) Weak-driving regime at low cost (*c* = 0.1, *κ* = 0); (C, D) strong-driving regime (*c* = 1, *κ* = 20). See Fig. S1 for approximate potentials in both regimes. Evolutionary parameters as in Fig. 2.

Importantly, however, code length changes do not happen alone, but they are always coupled to changes of the coding density. Moreover, given the difference in mutational clocks (*v* ≪ *μ*), the evolution of coding density settles to its stationary state before the next update of code length happens. These characteristics generate a fundamental asymmetry: code length extensions recruit new functional sequence units from random sequence, compression steps cut into existing functional sequence. Hence, extensions take place under net positive selection for recognition, compressions include a net constraint from locked-in target matches. These effects are described by average selection coefficients for extensions, *s*_+_(*ℓ*), and for compressions, *s*___(*ℓ*) (Fig. S2; see Methods for a precise definition of the averaging). Fig. 4BD displays these selection coefficients as arrows on the backbone of the evolutionary potential Φ(*ℓ*). This reveals an asymmetry that defines an evolutionary ratchet operating by selection: *s*_+_(*ℓ*) and *s*___(*ℓ* + 1) do not sum up to 0, and both selection coefficients differ from the increment of the potential. In other words, the ratchet clicks by different rules for forward and backward moves. The asymmetry of selection has a drastic consequence: the evolution of code length cannot be described by any fitness landscape that depends only on l. This would require *s*_+_(*ℓ*) = –*s*___(*ℓ* + 1) = *F*(*ℓ* + 1) – *F*(*ℓ*); the arrows in Fig. 4BD would collapse onto their backbone curve.

#### Weak driving, constraint ratchets

To capture the dynamical effects of ratchet evolution, we evaluate the total substitution rate for code length, 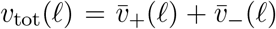, and the net elongation rate, 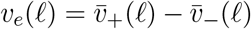. In the weak-driving regime, we find *v*_tot_ < 2*v*, indicating a net constraint on code length evolution (the ML rate 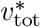 is shown in Fig. 3C, the length dependence in Fig. S2). At small *ℓ*, length extensions are approximately neutral and compressions are under constraint in functional sequence units (Fig. 4B), generating a net elongation rate *v_e_*(*ℓ*) ≲ *v* (Fig. S2). Hence, in this regime, an initially compact recognition site ratchets up slowly to a cost-dependent ML code length ℓ*. Notably, this length overshoots the point of maximal fitness defined by the landscape 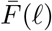 (Fig. 4A, Methods). Importantly, constraint ratchets produce complex recognition sites only at very low complexity cost, 2*N*_*c*_0__ ≪ 1 (Fig. 3AB). On larger scales of functional units, comparable ratchet mechanisms have been invoked to explain the complexity of RNA editing [7], cellular machines [8,10], and protein domains [11, 13–15].

#### Strong driving, adaptive ratchets

In the strong-driving regime, all length changes become adaptive (Fig. 4D), which generates fast ratchets with total click rates *v*_tot_(*ℓ*) and small-í elongation rates *v_e_*(*ℓ*) well above their neutral values (Fig. 3F, Fig. S2). In the analytical approximation, we find a universal ML click rate

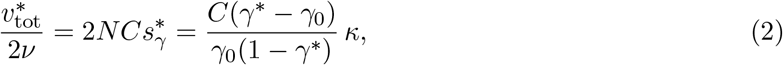

where *C* is a constant of order 1 (Methods). This shows that adaptive ratchets are driven by recognition target mutations. In contrast to constraint ratchets, adaptive ratchets generate molecular complexity also in the face of substantial costs (2*N*_*c*_0__ ~ 1). The ratchet dynamics of recognition sites has peculiar consequences for the evolution of recognition functions, to which we now turn.

#### Complexity increases the adaptive width

The most prominent effect of length increase is an increase in the susceptibility to selection. We measure this effect by the adaptive width A, defined as a suitably normalized expectation value of the population variance of the recognition trait, 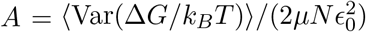. In the strong-driving regime and for a given recognition genotype, the adaptive width counts the mutational paths available for adaptive evolution of recognition; each path is weighted by its contribution to the speed of adaptation. Sites with larger A adapt their code faster at a given fitness gradient *∂F*/*∂*(Δ*G*); equivalently, such sites require less selection for recognition to maintain a given steady-state recognition *R* (Methods). The adaptive width is determined by the genetic architecture of the recognition trait; the actual speed of adaptation, or fitness flux, also depends on the global parameters *μ, N*, and on the fitness amplitude *f*_0_ [48]. In the minimal model, the ML adaptive width is related in a simple way to coding density and code length,

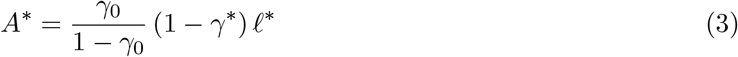

(Methods). In the strong-driving regime, *A*\* is a strongly increasing function of the driving rate (Fig. 3F). An increased adaptive width is clearly beneficial for maintaining function specifically in response to moving targets. By increasing molecular complexity, adaptive ratchets actually evolve increased values of A depending on the driving rate: the recognition system adapts its own adaptability to recognise moving targets.

#### Adaptive tinkering and functional alterations

Around the strong-driving ML point ℓ*, the ratchet dynamics of the code length becomes diffusive (|*v_e_*| ≪ *v*_tot_), as in near-neutral evolution, but the diffusion rate is adaptively enhanced (*v*_tot_ ≫ 2*v*, Fig. 3F). These characteristics define a ratchet regime of adaptive tinkering at large code length. The range of this regime is given by the width of the potential peak Φ(*ℓ*), which is defined as the r.m.s. variation Δ*ℓ* around ℓ* and can be computed from the curvature of the evolutionary potential. We find a broad regime of adaptive tinkering, Δℓ ~ ℓ* with a proportionality factor of order 1 (Fig. 4D, Methods). Rapid diffusion and broad length distributions are specific effects of adaptive ratchets. For evolution in a single-valued fitness landscape *F*(*ℓ*) = *F_r_*(*ℓ*) – *c*_0_ℓ, the diffusion rate at the peak is bounded by the neutral rate, 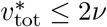. Moreover, if *F_r_* is a power of *ℓ*, length fluctuations around the peak scale as Δ*ℓ* ~ (ℓ*)^1/2^ (Methods).

Together, the turnover of recognition code includes point mutations at rates 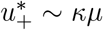 and length changes at rates 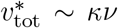, both of which are tuned by the seascape of target changes. Thus, by adaptive tinkering, recognition sites rapidly scan sequence space while their function is maintained. This enhances the probability of evolving functional refinements (for example, more specific target binding by epistatic interactions between sequence units) and of adopting new functions (examples are overlapping binding sites in bacteria [55, 56]). Evolutionary paths of recognition sites can also end with loss of function. Fig. S4 shows examples of such trajectories, where instabilities causing loss of function occur at different length scales. Short functional sites can be lost because they cannot keep up with a moving target of high *κ*; for longer sites, entropic forces and the cost of complexity can conspire to decrease Δ*G* and *ℓ*. We conclude that adaptive ratchets evolve a transient gain of molecular complexity, which can be locked in by additional functions or is exposed to pruning.

#### Complexity in immune recognition

The following example shows how selection on complexity can act in the adaptive immune system, a rapidly evolving recognition system of high global complexity [57]. In the presence of an antigen, immune B cells produce neutralizing antibodies that bind to specific target sites on its surface (called antigenic epitope sites) [58]. During and shortly after a primary infection, naive antibodies evolve high affinity to their cognate epitope by affinity maturation, a rapid evolutionary process under selection for recognition [16,59]. These antibodies are subsequently stored as immune memory to protect against future infections by the same pathogen. Fast-evolving pathogens, however, are moving targets: they change epitope sequences by accumulation of mutations, which can lead to eventual escape from immune recognition. This process, called antigenic drift, is frequently observed in RNA viruses, including human influenza and norovirus [60–62].

A minimal model of the recognition dynamics, sketched in Fig. 5A, suggests that fast antigenic drift can induce selection for a complex immune response. We assume a primary infection triggers the evolution of high-affinity antibody lineages targeting g distinct epitope sites of the pathogen (Δ*G_α_* = Δ*G_i_* > Δ*G*_50_ for *α* = 1,…,*g*). The breadth of the immune response, captured here by the parameter *g*, is known to vary substantially between viral pathogens and between human hosts [63,64]; it serves as a complexity measure for the immune response against a specific antigen [65]. Antigenic drift occurs by epitope mutations with effect ΔΔ*G*/*k_B_T* = *ϵ*_0_ and point substitution rate *ρ*; this process reduces the recognition *R_α_* of each epitope. Individual hosts are stochastically exposed to the evolving antigen with an average time *τ* between subsequent events [17]. An exposed individual gets infected if the antigen is not recognized at any of its epitopes, which occurs with probability Δ*R* = Δ*R*_1_ × ⋯ × Δ*R_g_*, where Δ*R_α_* = 1 – *R_α_*. This recognition function reflects the empirical finding that functional binding of a single epitope (monoclonal response) is sufficient to confer full protection to the host [63]. We then use a minimal fitness model of the form *F*(*g*) = *F_r_*(*g*) + *F_c_*(*g*), analogous to Eq. [1], for the host organism. Here, the recognition component *F_r_*(*g*) describes a given fitness cost per infection; the complexity component *F_c_*(*g*) accounts for the cost of evolving and maintaining epitope-specific antibody lineages, which we assume to be linear. Fig. 5B shows the expected steady-state fitness as a function of the number of epitopes for different values of the effective speed of antigenic evolution, *κ* = *ρτ*. We find a link between epitope complexity and target mutation rate remarkably similar to Figs. 4AC: multi-epitope immune responses provide a fitness benefit specifically against fast-evolving antigens (*κ* ≳ 1). A related effect has recently been discussed in ref. [27]: antigenic drift can favor generalist over specialist immune responses against multiple antigens. For even faster target evolution (*κ* ≫ 1), defence by immune memory becomes inefficient, in accordance with the results of ref. [17].

**Fig. 5:**
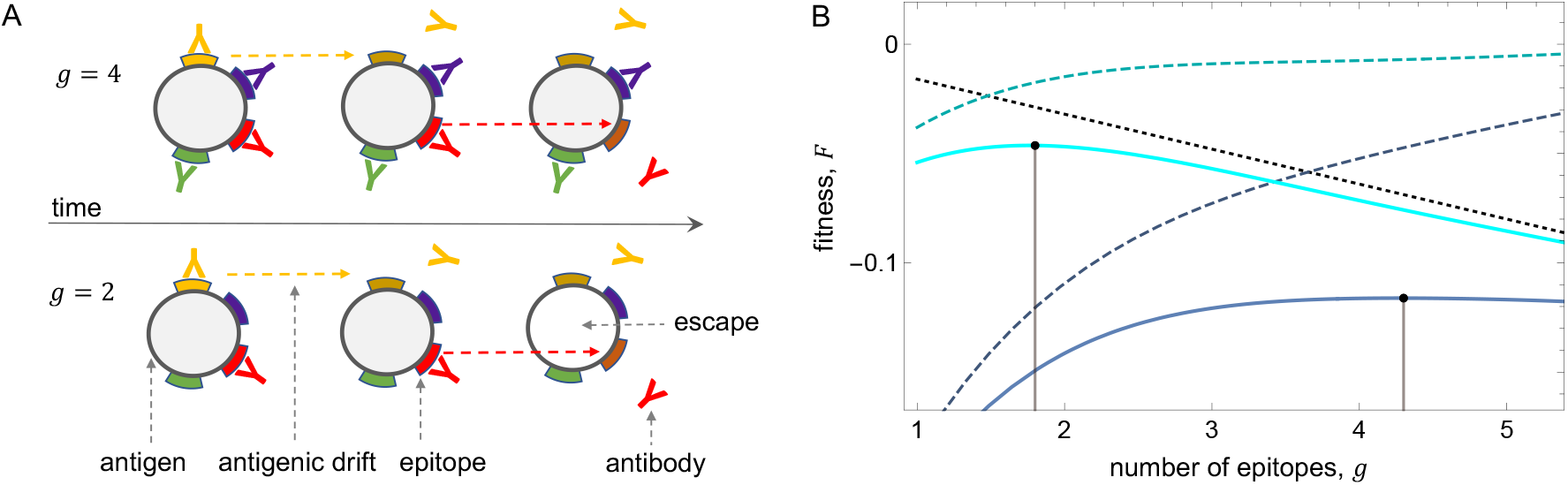
Complexity-dependent recognition and fitness of the adaptive immune system. (A) Immune recognition and escape evolution (schematic). Antibodies neutralize pathogens by binding to specific target sites (epitopes). The number of initially bound epitopes, *g*, measures the complexity of the immune response. Antigenic evolution of the pathogen (dashed arrows) changes the target sequences and reduces recognition in each epitope, eventually leading to escape from immune recognition. These dynamics is shown for *g* = 2 and *g* = 4. (B) Complexity-dependent fitness landscape, *F*(*g*) (solid), recognition component, *F_r_*(*g*) (dashed), and complexity cost, *F_c_*(*g*) (dotted black). ML values *g*\* are marked by black dots; *F_r_*(*g*) is plotted relative to the maximum recognition fitness, all fitness components are scaled by the exposure rate *τ*^-1^. This landscape is shown for slow- and fast-evolving pathogens with effective antigenic mutation rates *κ* = 2 (blue) and *κ* = 0.08 (cyan). To be compared with Fig. 4AC. Evolutionary parameters: *ϵ*_0_ = 2.5, Δ*G_i_* = Δ*G*_50_ + 2*ϵ*_0_, *f*_0_ = 1, *τc*_0_ = 0.016; the evolutionary algorithm is detailed in Methods.

This correlation between immune complexity and the speed of antigenic evolution may explain why multiple epitopes occur in several fast-evolving RNA viruses, including influenza [25,66], norovirus [62], and Sars-Cov-2 [67–69], although successful targeting of a single epitope is sufficient for neutralization [63]. Other features of immune recognition are not contained in the minimal model. These include a feedback of recognition onto the speed of target evolution. If epitope mutations come with a collateral cost to the pathogen (for example, through their effect on protein stability), a complex immune response can slow down or halt antigenic drift; this effect has recently been reported for measles [64]. The long-term evolution of epitope complexity, which depends on the density of naive antibody lineages in recognition space and on the affinity maturation machinery, is insufficiently understood at present and beyond the scope of this paper.

## Discussion

Organisms live in dynamic environments. Changing external signals continuously degrade the fidelity of an organism’s recognition units and generate selective pressure for change. Here we argue that stochastic, adaptive evolution of molecular interactions drives the complexity of the underlying sequence codes. Specifically, recognition sites tuned to a dynamic molecular target evolve a larger code length than the minimum length, or algorithmic complexity, required for function. The increase of coding length is coupled to a decrease of coding density; that is, individual letters carry a reduced information about the target. Importantly, these shifts also increase the adaptive width, defined as an importance-weighed number of adaptive paths for the recognition function, and the overall turnover rate of the recognition site sequence. Together, complex sites evolve in a specific regime of adaptive tinkering, which facilitates the maintenance of recognition interactions in dynamic environments. Moreover, the rapid parsing of fuzzy recognition site sequences increases the propensity for functional refinement or alteration, for example by tuning epistatic interactions within the recognition site or by recruiting additional targets for cooperative binding. We can summarize these dynamics as an evolutionary feedback loop: moving targets, by increasing the molecular complexity of their cognate recognition machinery, improve their own recognition.

Our analysis suggests specific ratchet modes of evolution by which recognition sites gain molecular complexity. All evolutionary ratchets acting on sequence code build on a fundamental asymmetry: code length extensions take place into random sequence, compression steps cut into functional sequence. Thus, even without any explicit fitness benefit, complexity can evolve as a collateral of selection for function. If the cost of complexity is small, evolutionary ratchets can operate by neutral steps of complexity increase. In Lynch and Conery’s model of macro-evolution, the genomes of higher organisms grow neutrally because reduced effective population sizes reduce the cost of complexity; later, functions can latch on and eventually be conserved by selective constraint [9]. Similarly, in entrenchment models of protein evolution, complex multi-domain states establish neutrally if the cost of formation is small. Such complexes can be maintained by an accumulation of mutations that are conditionally neutral but deleterious in simpler states [14]. In contrast, adaptive ratchets generate positive selection on complexity changes coupled to functional adaptation; this mechanism can override the cost of complexity. The rate of adaptive ratchets is set by external driving forces, here recognition target changes, not by genetic drift. Hence, unlike classical arguments of complexity evolution, adaptive ratchets do not describe the refinement of functions in a static environment but the maintenance of functions in the face of fitness seascapes.

How universal are evolutionary ratchets? We can expect this mechanism to operate in a broad range of molecular networks with multiple parallel sub-units and a reservoir of potential units. Examples include recognition sites with multiple sequence matches, repertoires of immune response with multiple antibody lineages, and on larger scales, regulatory or metabolic networks with multiple genes contributing to the same function. Such networks allow ratchet modes of evolution to kick in: by preferential attachment of fit and deletion of unfit sub-units, changes in network architecture can couple to selection for network function. Specifically, the adaptive ratchet model then predicts that external driving – here by moving recognition targets, evolving antigens, or time-dependent metabolic substrates – induces the evolution of complex networks with redundant sub-units.

New high-throughput methods are currently used to map sequence and phenotypes of molecular interactions at an unprecedented depth. Prominent examples include transcription factor interactions in microbes [22,70,71] and receptor-antigen interactions in human immune repertoires [72–74]. This opens the perspective of new experimental screens to map correlations between external driving, network architecture, and molecular complexity. Experiments with exposure to moving targets can determine the dominant selective force – constraint or adaptation – driving the ratchet evolution in these systems. Such experiments can also become a powerful tool of evolutionary control, to elicit molecular complexity of recognition and its potential for evolutionary innovation.

## Acknowledgments

We thank Rob Phillips for discussions. This work was supported by Deutsche Forschungsgemeinschaft grant CRC 1310 “Predictability in Evolution”.

## Code Availability

All code used to run simulations and produce figures is available in a Github repository: https://github.com/tomroesch/adaptive_ratchets

## Methods

### Recognition trait and function

In a biophysical model of recognition, a quantitative trait characterising a recognition site is its excess binding affinity to the recognition target, Δ*G*, compared to random sequence loci. Here we use a minimal model with a reduced affinity

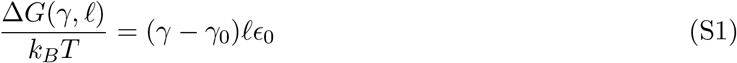

at physiological temperature *T*, which depends on the site length, *ℓ*, the fraction of target-matching sequence units in the site, *γ* = *k/ℓ*, the average fraction of matches in random sequence, *γ*_0_, and the reduced affinity gain per match, ΔΔ*G*/*k_B_T* = *ϵ*_0_. The recognition function is defined as the equilibrium target binding probability,

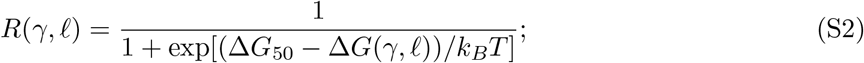

this function depends on Δ*G*/*k_B_T* and the half-binding point under physiological conditions, Δ*G*_50_/*k_B_T* = ℓ_0ϵ0_.

### Sequence evolution rates

In the minimal model, the sequence evolution of recognition involves the three classes of changes shown in Fig. 1. (i) Point mutations of the recognition sequence change the coding density at constant code length and occur at a rate *μ* per unit of sequence (Fig. 1A). These changes have point mutation rates

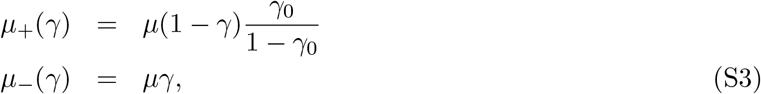

and selection coefficients

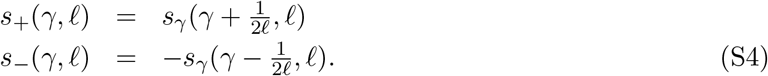

depending on the direction of selection (beneficial/deleterious, indicated by the index ±). Here

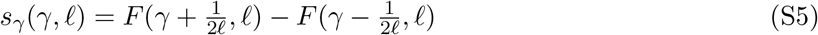

is the fitness increment in the landscape (1) in midpoint discretisation. Eqs. (S3) and (S4) determine the substitution rates

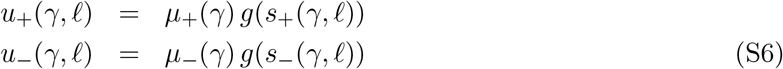

in the low-mutation regime (*μ*N*ℓ* ≲ 1). Here *g*(*s*) = *Nπ*(*s*) is Haldane’s factor in a population of effective size *N* [75],

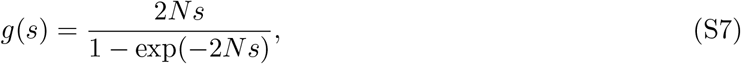

which is proportional to the fixation probability of an individual mutation, *π*(*s*). With an appropriately changed effective population size, these substitution rates apply even if the global sequence evolution takes place under clonal interference [54,76]. (ii) Recognition site extensions and compressions add or remove one unit of sequence (Fig. 1BC) and occur at a rate *v* ≪ *μ*. These changes have mutation rates

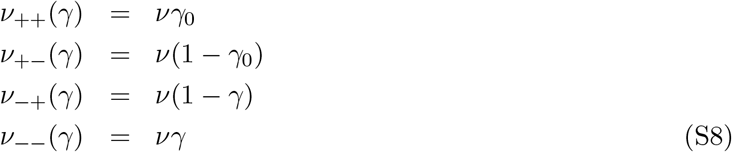

and selection coefficients

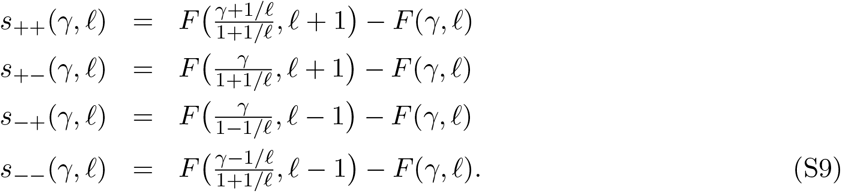

Using the exponential approximation of the target binding probability

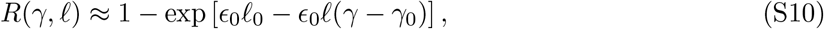

we can relate these selection coefficients to the selection coefficient for point mutations of the recognition sequence in Eq. (S5),

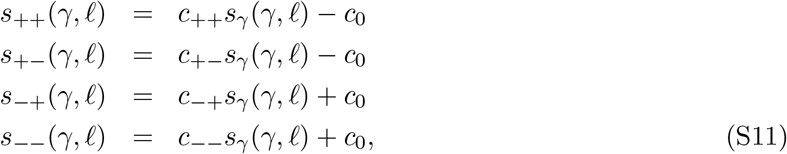

where *c*_0_ denotes the complexity cost parameter in the fitness landscape [1] and

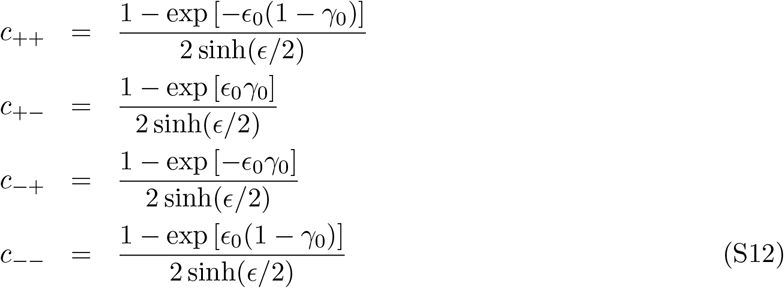

Here we compute one selection coefficient explicitly,

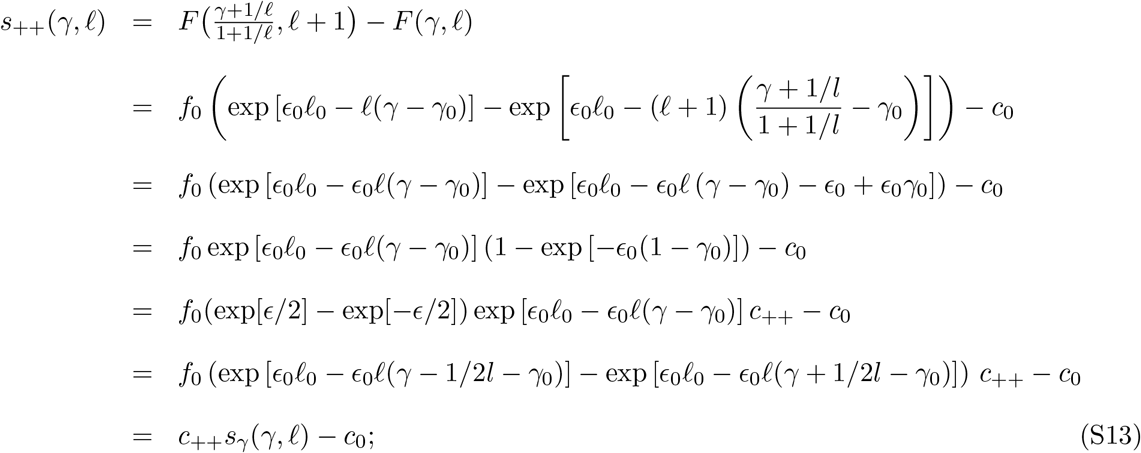

the other coefficients are obtained in the same way. These selection coefficients depend on the direction of change (extension/compression, first index ±) and the direction of selection (beneficial/deleterious, second index *±*). The approximate expressions in terms of the fitness increment *s_γ_* are used for the analytical closure discussed below. The exponential prefactors, which are of order 1, reflect the shift in the half-binding point *γ*_50_ induced by a length change and the switch to the midpoint discretization in Eq. (S5). Eqs. (S8) and (S9) determine the total substitution rates of extensions and compressions,

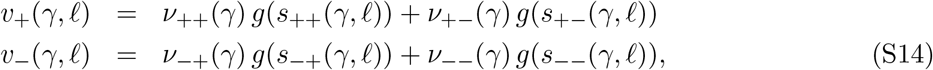

(iii) Recognition target changes occur at a rate *ρ* = *κμ* per unit of sequence, independently of the recognition function (Fig. 1D). Hence, beneficial and deleterious target changes have rates

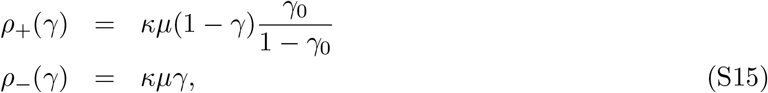

which have the same combinatorial factors as the point mutations (S3) and are independent of the fitness landscape *F*(*γ,ℓ*).

Individual evolutionary trajectories (*k,ℓ*)(*t*) generated by the substitution dynamics with rates *u*_±_(*γ,ℓ*), *v*_±_(*γ, ℓ*), and *ρ*_±_(*γ*) are shown in Fig. 2BD and Fig. S4.

### Evolutionary potentials and stationary distribution

Given the separation of mutational time scales for coding density and code length (*v* ≪ *μ*), the stationary distribution factorizes,

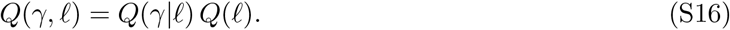

The conditional distribution of the coding density takes the form

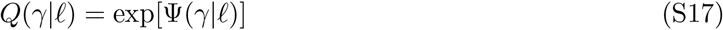

with a potential function given by detailed balance,

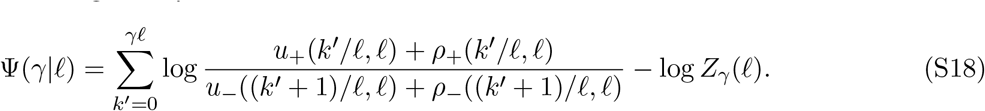

Here *Z_γ_*(*ℓ*) is a normalization factor and the point substitution rates *u*_±_, *ρ*_±_ are given by Eqs. (S6) and (S15). This distribution defines the conditional average of an observable *x*(*γ, ℓ*),

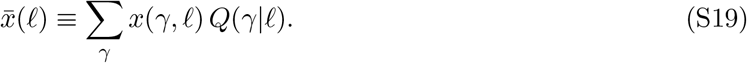

Because *Q*(*γ*|*ℓ*) is approximately Gaussian (Fig. S1), such averages can be accurately evaluated by steepest descent, 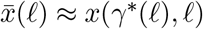, where *γ*\*(*ℓ*) = argmax_*γ*_ *Q*(*γ*|*ℓ*) denotes the conditional ML coding density. In particular, this procedure yields an effective fitness landscape for code length,

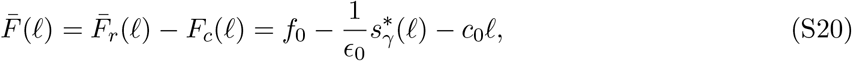

where 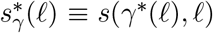 is the selection coefficient at the ML coding density. Here we use an exponential approximation of the Hill recognition function *R*(*γ,ℓ*) in the regime of functional sites, which implies 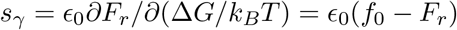. The marginal distribution of code length,

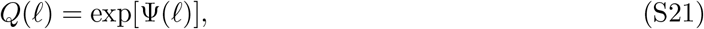

is again determined by detailed balance,

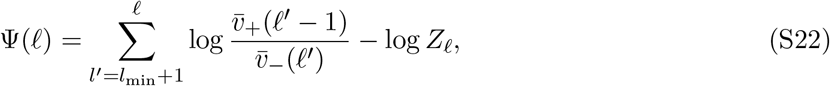

where *Z_ℓ_* is a normalization factor and 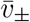 are the average extension and deletion rates given by Eqs. (S14) and (S19). From the full distribution *Q*(*γ,ℓ*), we define the corresponding distribution for functional sites,

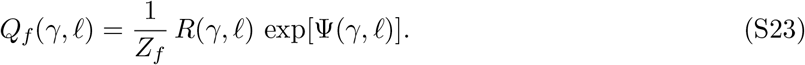

This distribution weighes sequence states by the recognition (target binding) probability, *R*(*γ,ℓ*). Examples of the full distribution *Q*(*γ*|*ℓ*) = exp[Φ(*γ*|*ℓ*)] are plotted in Fig. 2AC; the evolutionary potential Φ(*ℓ*) is shown in Fig. 4BD.

### Weak-driving regime

For *κ* ≲ 1, the evolutionary potential for coding density takes the form

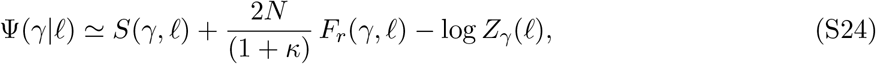

where

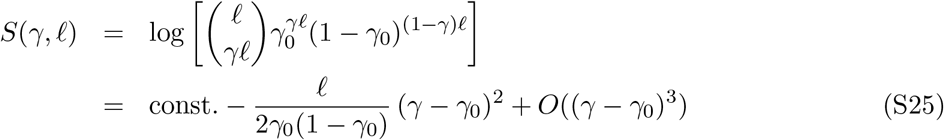

is the entropy, defined as the log number of sequence states, at a given recognition density. At equilibrium (*κ* = 0), this potential reduces to the well-known free fitness that generating the Boltzmann-Gibbs distribution of the substitution process (S6) [36,50]. The extension to driven processes can be derived a weak-selection approximation, which is appropriate close to equilibrium. In this regime, we expand the rates *u*_±_(*γ, ℓ*) and *ρ*_±_(*γ*) to first order in the selection coefficient *s_γ_*(*γ,ℓ*). We can then write the stochastic dynamics of the mean coding density *γ* as a diffusion process [54,77],

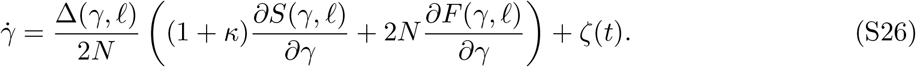

The stochastic force *ζ*(*ℓ*) has mean 〈*ζ*(*ℓ*)〉 = 0 and covariance

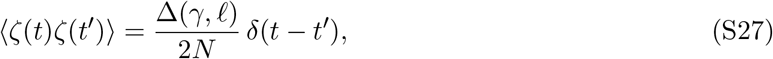

which is related to the diversity of the coding density in a population of size *N*,

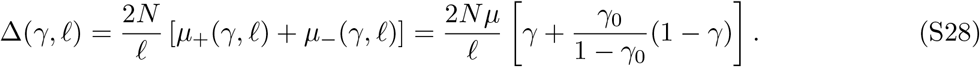

The neutral process includes recognition site and target mutations, leaving the neutral distribution *Q*_0_(*γ*|*ℓ*) = exp[*S*(*γ,ℓ*)] unaffected by the driving. In contrast, the population diversity Δ includes only site mutations. This diversity governs the response to selection in Eq. (S26), while target mutations are independent of the fitness landscape *F*(*γ,ℓ*). Maximizing the potential Φ(*γ*|*ℓ*) determines the conditional ML point γ*(*ℓ*). This point satisfies a balance condition between selection, site mutations and the associated genetic drift, and target mutations. At equilibrium, this reduces to the well-known detailed balance between deleterious and beneficial substitutions (Fig. 2B). By Eqs. (S24) and (S25), the weak-coupling balance condition relates selection and coding density,

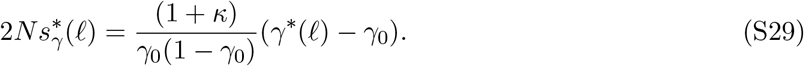

The evolutionary potential for length takes the weak-driving form

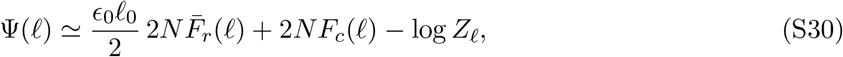

where *ℓ*_0_ = Δ*G*_50_/(*ϵ*_0_*k_B_T*) and the recognition component, 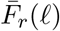, is determined by Eq. (S20). To derive this form, we use again a weak-selection approximation and expand the substitution rates for extensions and compressions, *v*_±_(*γ,ℓ*), to first order in the selection coefficients *s_γ_* and *c*_0_. From Eqs. (S8)–(S14), we obtain

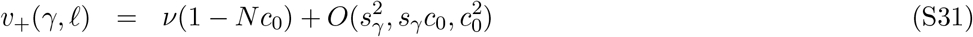

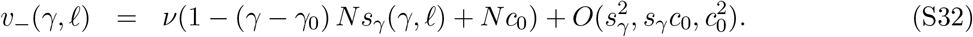

We can now compute the potential increment in the steepest-descent approximation for *γ*,

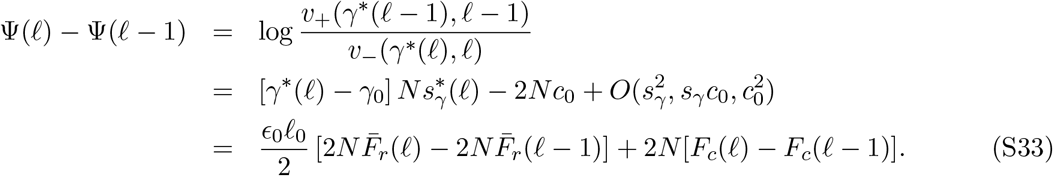

The last approximation uses that selection is proportional to the recognition load, *s_γ_* = *ϵ*_0_(*f*_0_ – *F_r_*), and its ML value is approximately inversely proportional to the site length, 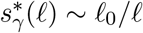, by Eqs. (2) and (S29). Eq. (S30) then follows by summation of the increment up to a given value of *ℓ*. Maximizing Φ(*ℓ*) determines the balance condition for code length,

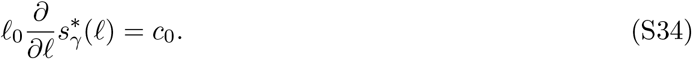

Together with Eq. (S29) and the leading-order relation between coding density and length, Eq. (2), we obtain an analytical closure for the global ML point (*γ*,ℓ\**) in the weak-driving regime,

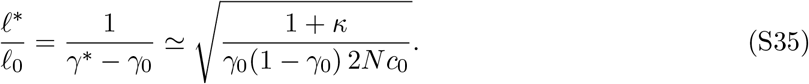

Compared to the effective fitness 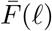, as given by Eq. (S20), the potential Φ(*ℓ*) upweighs the recognition component, which implies that the global ML length *ℓ*\* overshoots the maximum of 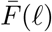 in the weak-driving regime (Fig. 4A). In Fig. S1AB, we compare the full potentials Φ(*γ|ℓ*) and Φ(*ℓ*) with their the weak-coupling approximations; Fig. S3A shows that this approximation produces the correct scaling of the equilibrium ML point with the cost parameter *c*_0_. In particular, the condition that evolutionary equilibrium leads to near-maximal coding density, *γ*\*(*κ* = 0) ≈ 1, sets a constraint roughly equal to the selection for coding density, 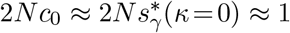. This informs our choice of the cost parameter for the crossover crossover to strong driving.

### Strong-driving regime

For *κ* ≳ 1, selection on code changes becomes strong (*s*\*(*ℓ*) ≫ 1), causing deleterious point mutations to freeze out. The resulting balance condition for the ML coding density involves beneficial site mutations and target mutations (Fig. 2D),

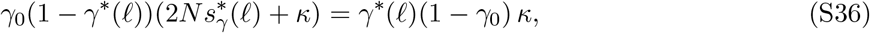

Like its weak-selection counterpart, Eq. (S29), this condition relates selection and coding density,

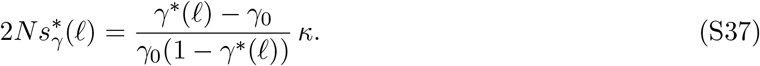

Similarly, strong selection causes deleterious extensions and compressions to freeze out (2*N*_*s*+__ ≪ 1, 2*Ns*_− −_ ≪ 1), and the balance condition for code length becomes

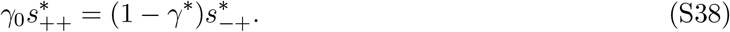

Inserting the approximations (S9) and solving for 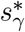, we obtain

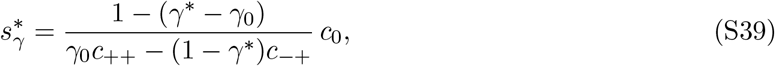

which is the strong-selection counterpart of Eq. (S34). The analytical closure for the global ML point in the strong-driving regime given by Eqs. (S37) and (S39),

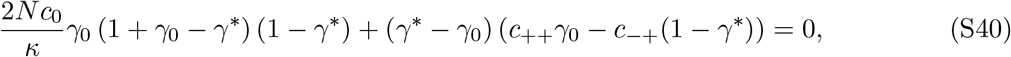

determines the ML point (*γ*,s\**); the ML code length, *ℓ*\*, is then obtained from Eq. (2). The full potentials Φ(*γ|ℓ*) and Φ(*ℓ*) and their strong-coupling approximations are shown in Fig. S1. The ML closure accurately captures the crossover to the strong-driving regime (Fig. S3).

### Ratchet evolution

The ratchet dynamics derived in this paper is defined by the breakdown of detailed balance for evolution under selection. Specifically, the minimal model for recognition sites generates net selection coefficients for code length changes,

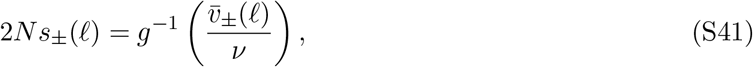

where *v* is the basic mutation rate of extensions and compressions, 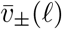 are the corresponding substitution rates defined by Eqs. (S14) and (S19) (Fig. S2). The function *g*^−1^ is the inverse of the Haldane function *g* defined in Eq. (S7). Ratchet evolution of code length is defined by a decoupling of selection for reverse changes,

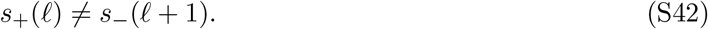

This inequality says that the substitution rates 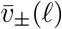 cannot be generated from the neutral rate *v* and selection by any fitness landscape *F*(*ℓ*), which would imply *s*_+_(*ℓ*) = –*s*___(*ℓ* + 1) = *F*(*ℓ* + 1) – *F*(*ℓ*) and a detailed balance relation of the form *v*_+_(*ℓ*)/*v*___(*ℓ* + 1) = *g*(*s*+(*ℓ*))/*g*(*s*___(*ℓ* + 1)) = exp[2*N*(*F*(*ℓ* + 1) – *F*(*ℓ*)]. In particular, the selection coefficients *s*_±_(*ℓ*) decouple from the increment of the effective fitness landscape for code length, 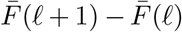, as defined by Eq. (S20) (Fig. 4AC). This landscape maps the average long-term fitness effects of length changes but does not govern the ratchet dynamics. The selection coefficients *s*_±_(*ℓ*) also decouple from the increment of the evolutionary potential, Φ(*ℓ* + 1) – Φ(*ℓ*) (Fig. 4BD). In the stationary state of the ratchet, the substitution rates satisfy the detailed balance relation 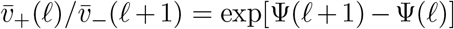, but the link of these rates to the neutral rate *v* by Haldane’s formula is lost.

### Adaptive tinkering

The ratchet substitution dynamics of extensions and compressions can be characterized by the total rate 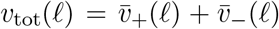, which sets the overall speed of code length evolution, and the net elongation rate, 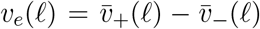 (Fig. S2). Inspection of the potential Φ(*ℓ*), Eq. (S22) with rates given by Eq. (S14), shows that the total rate takes the form 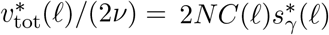 with a slowly varying coefficient function *C*(*ℓ*) ~ 1. In particular, we obtain an analytical estimate for the total ratchet rate at the ML point, as given by Eq. (3). This relation implies 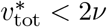 in the weak-driving regime, where 2*Ns_γ_* ~ 1, and *v*_tot_ > 2*v* for *κ* ≫ 1, in accordance with the evaluation from the full rates (Fig. 3CF, Fig. S2).

In the strong-driving regime, the range of adaptive tinkering around the ML point can be defined by the r.m.s. code length fluctuations in the stationary state,

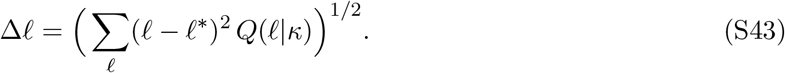

We estimate Δ*ℓ* from the curvature of the evolutionary potential for length,

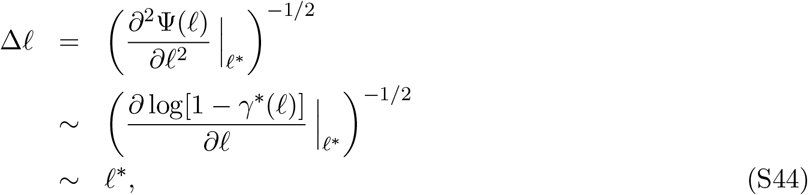

To derive this scaling, we Taylor-expand the potential Φ(*ℓ*) around the ML point *ℓ*\*, using Eq. (S22) with rates given by Eq. (S14) in the strong-driving regime (2*Ns*_+__, 2*Ns*_− −_ ≪ 1). Then we use the leading-order scaling *γ*\* (*ℓ*) – *γ*_0_ ~ *ℓ*^−1^, consistent with Eq. (2). The analytical approximation of Eq. (S44) is in accordance with the tinkering range evaluated from the full distribution *Q*(*ℓ*) (Fig. 3E). The ratchet generates anomalously broad length fluctuations compared to evolution in a uni-valued fitness peak of the form *F*(*ℓ*) = *f*_0_ – *c_r_ℓ*^−*α*^ – *c*_0_*ℓ* (*α* > 0). In that case, the stationary-state fluctuations scale differently, Δ*ℓ* ~ (*ℓ*\*)^1/2^, as can be seen by evaluating the Boltznann-Gjibbs distribution *Q*_eq_(*ℓ*) exp[2*NF* (*ℓ*)].

### Adaptive width and fitness flux

We define the adaptive width of a quantitative trait, here Δ*G*, as its normalized heritable variation,

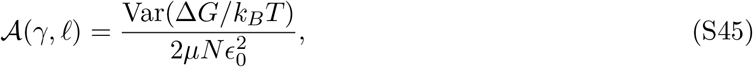

where 2*μN* is the neutral sequence diversity per unit of length and 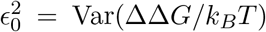 is the variance of the mutational effect distribution. The adaptive width is a measure of susceptibility to selection [78]. This can be seen by relating it to the change in mean population fitness by natural selection, 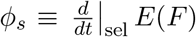, in a trait-dependent fitness landscape *F*(Δ*G*). By Fisher’s theorem, we obtain

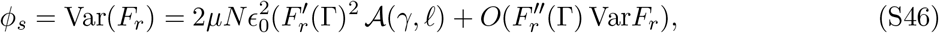

where Γ = *E*(Δ*G*/*k_B_T*) denotes the population mean trait. The selective flux *ϕ_s_* is a non-negative component of the fitness flux *ϕ*. In an evolutionary steady state, *ϕ_s_* measures the amount of selective turnover required to maintain a time-independent average trait 〈Γ〉. The total fitness flux *ϕ* also includes the fitness change by evolution under mutations, selection, and genetic drift [48]. It vanishes in an evolutionary equilibrium state. In a driven stationary state, it has a time-independent average, 〈*ϕ*〉 > 0, that balances with the rate of fitness change by external fluctuations.

In a model with site-dependent trait effects *ϵ_i_*(*i* = 1,…,*ℓ*), we obtain a normalized trait diversity

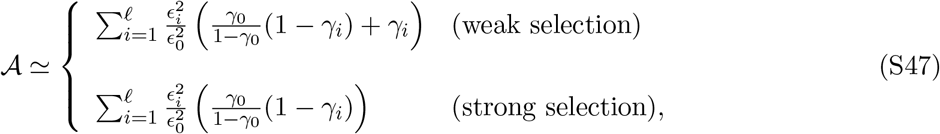

where *γ_i_* = 1 for site positions with a target match and *γ_i_* = 0 for positions with a mismatch. The weighting factors 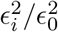 reflect the higher contributions of strong-effect mutations to the adaptive process; different regimes for transient adaptive dynamics have been discussed in ref. [79]. In the weak-selection regime 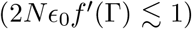, all positions contribute an expected sequence diversity 2Nμ, generating the maximum trait diversity [77,78]. In the strong-driving regime (2*Nϵ*_0_*f*′(Γ) ≫ 1), the trait diversity narrows to positions with deleterious majority alleles (*γ_i_* = 0). These positions contribute an expected trait diversity 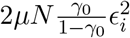, predominantly from mutations destined for fixation. Positions with beneficial majority alleles (*γ_i_* = 1) freeze out; these positions have a much smaller trait diversity 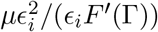. Together, the adaptive width 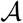 depends on the distribution of effects *ϵ_i_* and the occupancy of beneficial and deleterious sequence states but decouples from the global parameters *μ, N*, and from the overall strength of selection, *F*′(Γ).

Here we focus on the expectation value of the adaptive width over parallel-evolving populations, 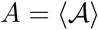. In the strong-selection regime of the minimal model (*ϵ*_*i*_ = *ϵ*_0_, 〈*γ_i_*〉 = *γ*), we obtain

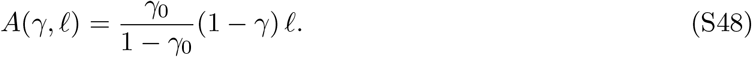

At the global ML point (*γ*\*,*ℓ*\*)(*κ*), this relation reduces to Eq. (4); the function *A*\*(*κ*) is shown in Fig. 4F. The adaptive width can be compared to the expected adaptive speed of the recognition trait by point substitutions,

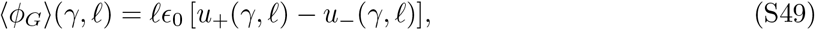

and the corresponding fitness flux,

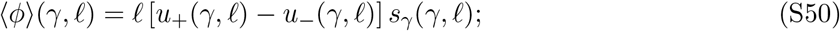

the contribution of length changes to the adaptive fluxes is down by a factor *O*(*ν/μ*). In the strong-driving regime, we obtain

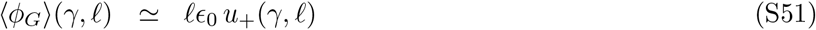

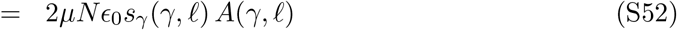

and

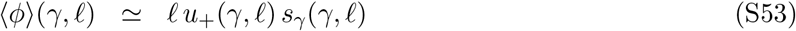

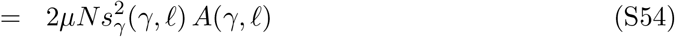

using Eq. (S6). Thus, at a given value of *s_γ_*, the expected fluxes 〈*ϕ_G_*〉 and 〈*ϕ*〉 increase with *A*, while the selection *s_γ_* required for a given value of 〈ϕ_G_〉 decreases with increasing *A*. In an evolutionary steady state, the adaptive fluxes match the effect of recognition target changes, as given by the strong-driving balance condition (S37). We obtain

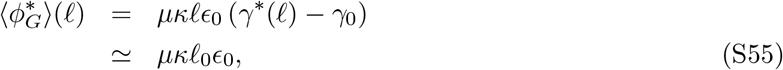

and

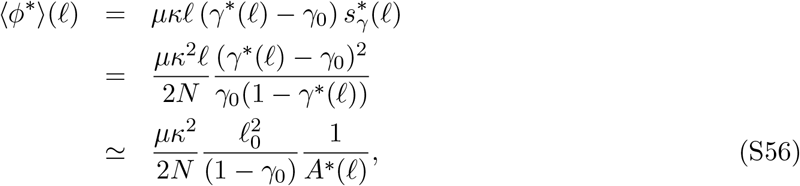

where we we use the leading-order scaling *γ*\*(*ℓ*) – *γ*_0_ ~ *ℓ*^−1^, consistent with Eq. (2). According to these relations, the adaptive speed of the recognition trait becomes asymptotically independent of *ℓ*; the fitness flux required to maintain the stationary state decreases with increasing adaptive width.

### Evolution of immune recognition

Our minimal dynamics model has three characteristics: (i) Infections by a given pathogen (antigen) induce the affinity maturation of antibodies targeting *g* distinct epitopes. Functional binding, leading to neutralization of the antigen, is described by a Hill recognition function, as given by Eq. (S2). The antibodies produced by an infection at time *t*_0_ are assigned an initial affinity Δ*G_α_*(*t*_0_) = Δ*G_i_* = Δ*G*_50_ + 2*ϵ*_0_ to their cognate epitopes (*α* = 1,…,*g*). (ii) Antigenic drift is a Poisson process that changes epitope sequences with a point substitution rate *ρ*; each epitope mutation has a reduced affinity effect *ϵ*_0_ = ΔΔ*G*/*k_B_T*. Antigenic drift generates time-dependent affinities Δ*G_α_*(*t*) (*α* = 1,…,*g*). (iii) Exposure of an individual to the evolving antigen is an independent Poisson process with rate *τ*^−1^. For an exposure event at time *t*, we evaluate the recognition function

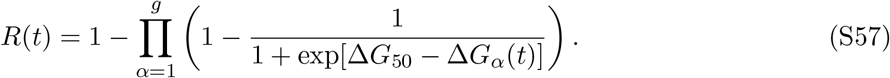

Neutralization by activation of immune memory occurs with probability *R*(*t*), assuming that each epitope is an independent and sufficient channel for recognition of the antigen. A full secondary infection occurs with probability Δ*R*(*t*) = 1 – *R*(*t*), resetting all affinities to Δ*G*_0_. Here we assume that significant updating of the immune response by affinity maturation occurs only after full infections. Additionally, exposure events that occur between full infection events can boost antibody titers against the corresponding pathogens unevenly [80]. However, this does not affect our results that rely on the underlying immunological memory rather than on sera antibody titers.

We simulate long-term affinity trajectories (Δ*G*_1_,…, Δ*G_g_*)(*t*) for the joint dynamics of antigenic drift and exposures. The effective speed of antigenic evolution, *κ* ≡ *ρτ*, is given by the expected number of mutations per epitope between subsequent exposures. We record the steady-state average 〈Δ*R*〉(*g, κ*); this quantity is proportional to the fitness cost of infections and enters a minimal host fitness model of the form of Eq. (1), where the cost of complexity is assumed to be linear with slope *c*_0_. The resulting fitness landscape *F*(*g*|*κ*) determines the optimal epitope complexity for given antigenic speed, *g*\*(*κ*). For low to moderate antigenic drift (*κ* ≲ 2), *g*\*(*κ*) increases with *κ* (Fig. 5B). For higher values of *κ*, the number of epitopes needed for protection increases exponentially, inducing a prohibitive cost and a transition to *g*\*(*κ*) = 0. A negative correlation between the speed of antigenic evolution and the efficacy immune memory has also been mapped in multi-pathogen models [17].

### Numerical evaluation

Code for evaluation of full and approximate analytical expressions (Fig. 2 – 4, S1 – S3), and for numerical simulations of evolutionary trajectories (Fig. 2, 5, S4) is provided as Supporting Information.

## Supporting Figures

**Fig. S1:**
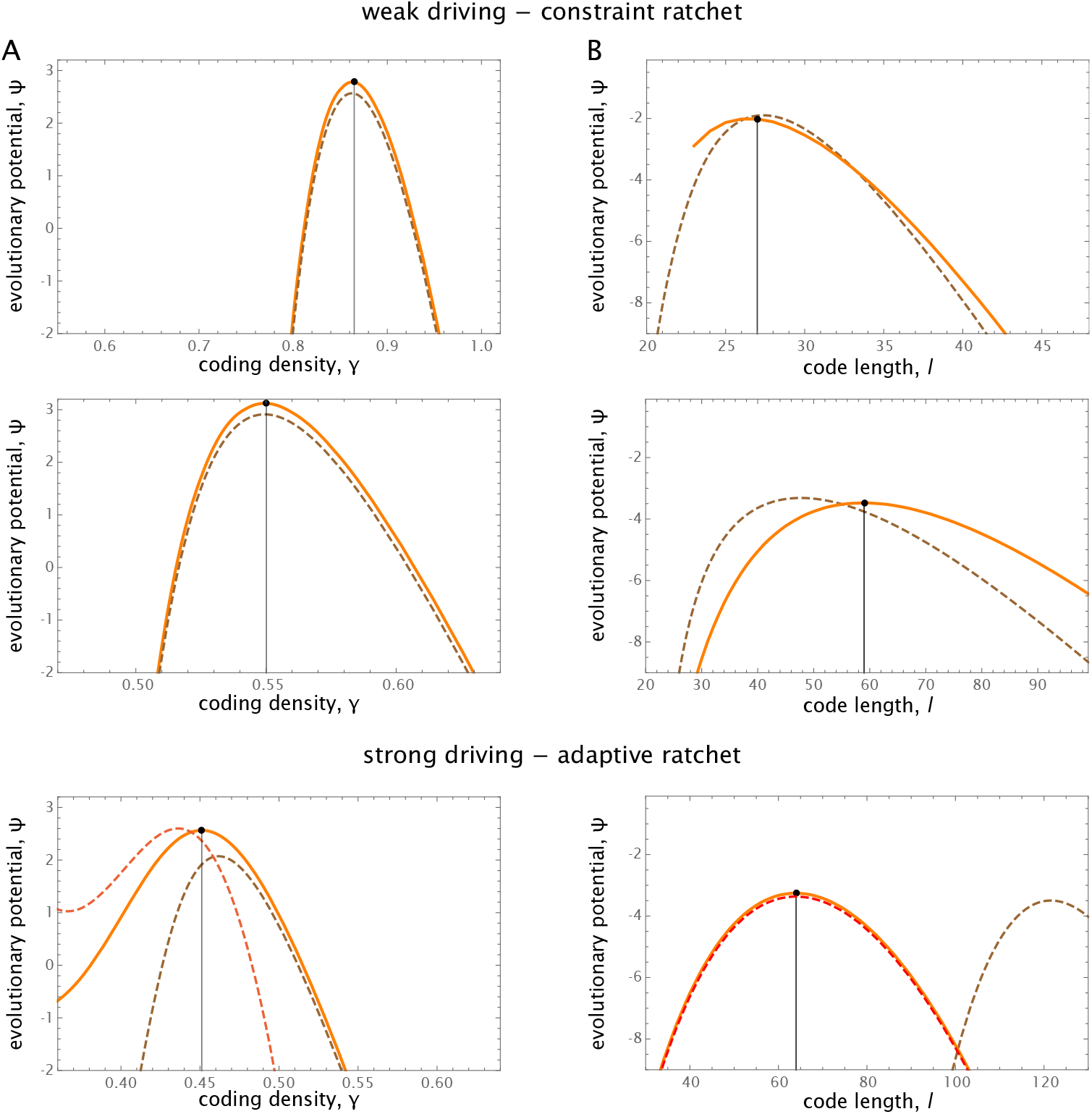
Approximate evolutionary potentials. (A) Evolutionary potential for coding density at ML code length, Ψ(*γ|ℓ*\*). (B) Evolutionary potential for code length, Ψ(*ℓ*). Full potentials (orange) are compared to the corresponding weak-driving (brown) and strong-driving (red) approximations. Parameters: (top) weak-driving regime, 2*N*_*c*_0__ = 1, *κ* = 0; (center) weak-driving regime at low cost, 2*N*_*c*_0__ = 0.2, *κ* = 0; (bottom) strong-driving regime, 2*N*_*c*_0__ = 1, *κ* = 20. Other parameters as in Fig. 2.

**Fig. S2:**
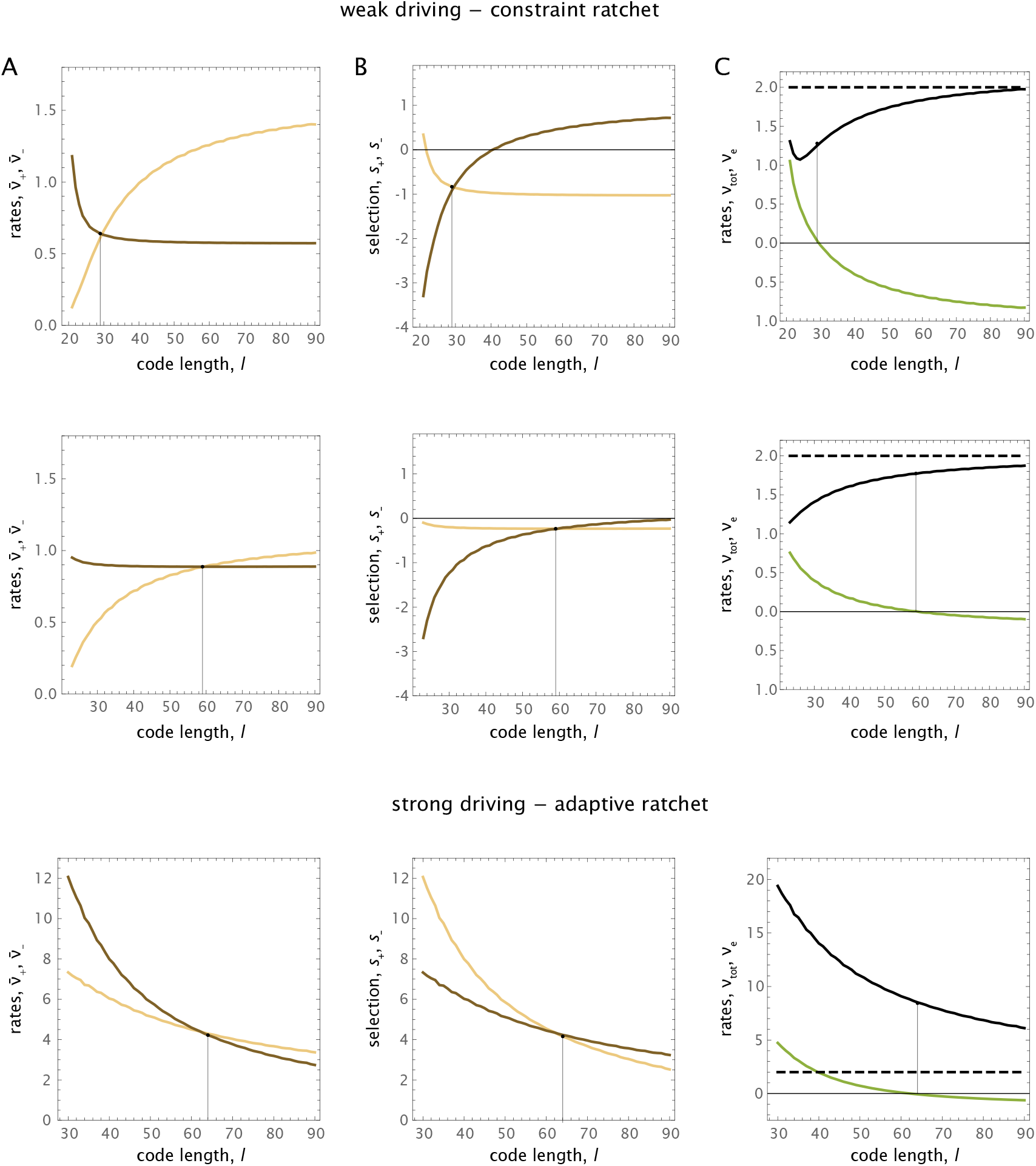
Ratchet dynamics. (A) Substitution rates for code extension and compression, 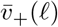 (light) and 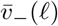 (dark). (B) Ratchet selection coefficients for extension and compression, *s*_+_(*ℓ*) (light) and *s*___(*ℓ*) (dark). (C) Total ratchet rate, *v*_tot_(*ℓ*) (black), and net elongation rate, *v_e_*(*ℓ*) (green). Parameters: (top) weak-driving regime, 2*N*_*c*_0__ = 1, *κ* = 0; (center) weak-driving regime at low cost, 2*N*_*c*_0__ = 0.2, *κ* = 0; (bottom) strong-driving regime, 2*N*_*c*_0__ = 1, *κ* = 20. Other parameters as in Fig. 2.

**Fig. S3:**
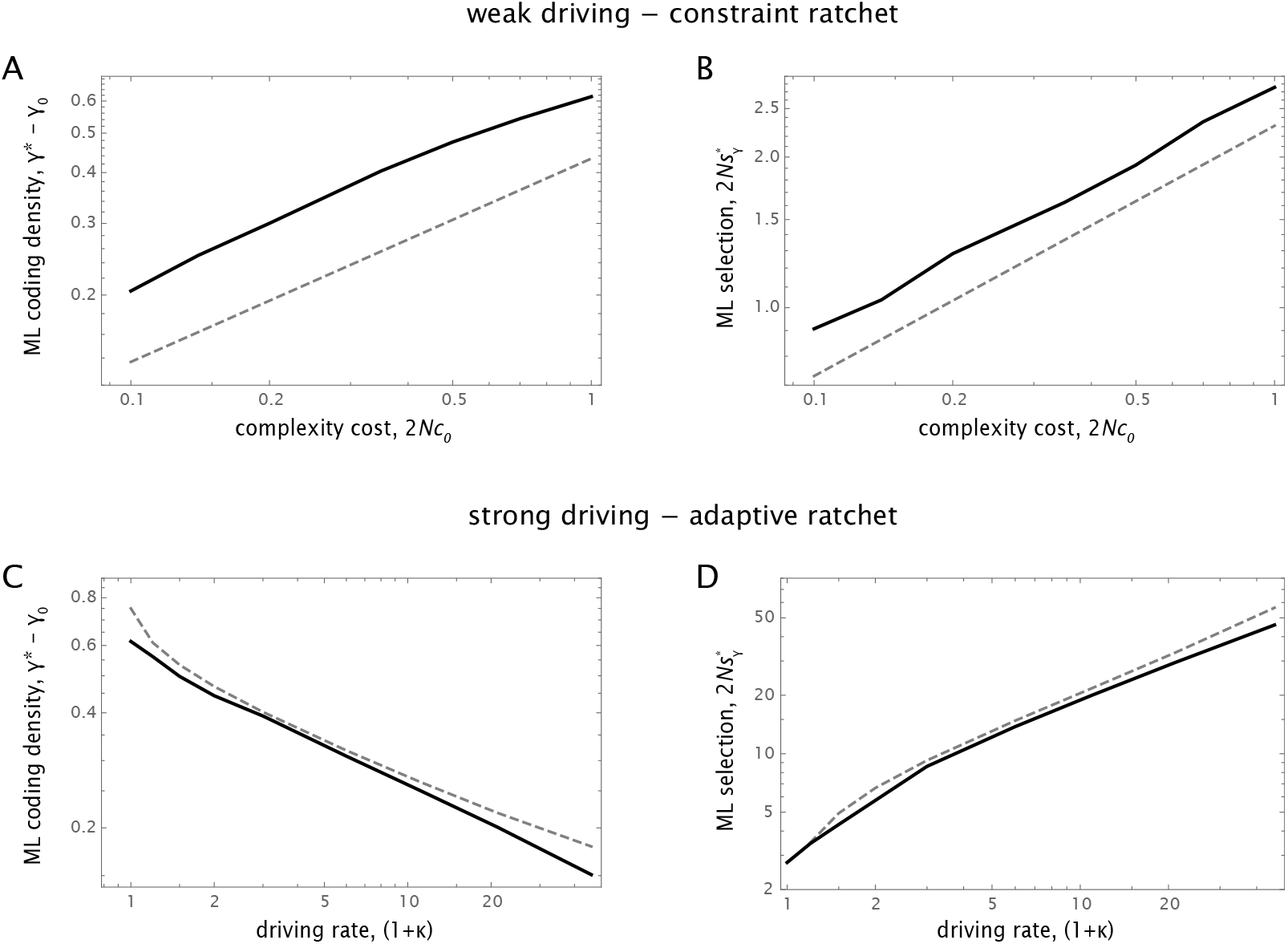
Global ML likelihood points. (A, B) Crossover to low cost. Full solution (solid) and weak-coupling approximation (dashed) for coding density, γ*(2*N*_*c*_0__, *κ* = 0) and selection on recognition, 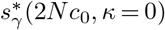. (C, D) Crossover to strong driving. Full solution (solid) and strong-coupling approximation (dashed) for coding density, γ*(2*N*_*c*_0__ = 1,*κ*) and selection on recognition, 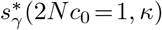. Other parameters as in Fig. 2.

**Fig. S4:**
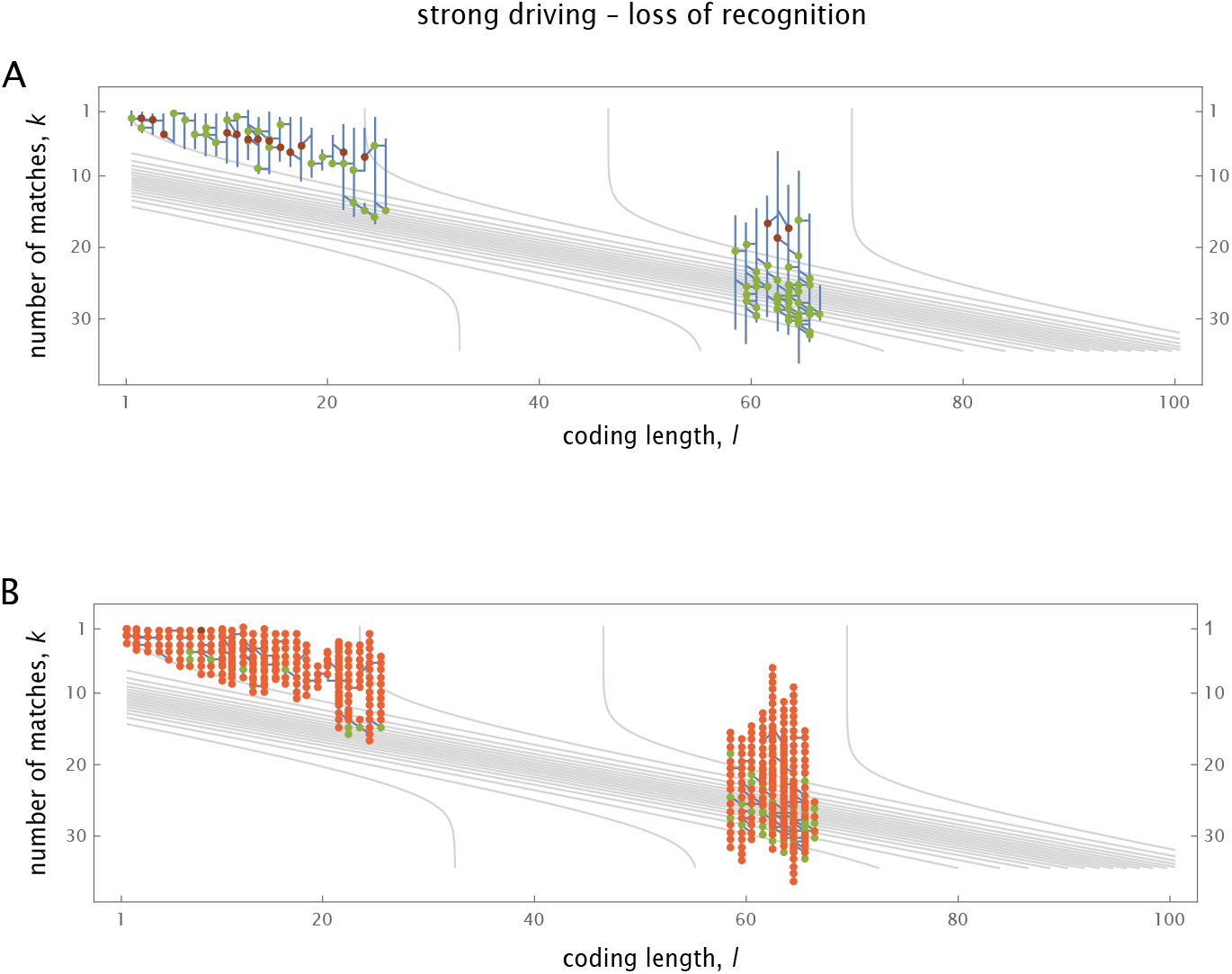
Evolutionary paths with loss of function. Loss of function by degradation at low code length (left, *c* =1, *κ* = 10, 10000 mutational time steps, starting at *ℓ* = 20) and at high code length (right, 2*N*_*c*_0__ = 1, *κ* = 45, 2000 mutational time steps, starting at *ℓ* = 60). (A) Coding density changes (green: adaptive recognition site mutations, red: recognition target mutations). (B) Code length changes (green: adaptive, purple: deleterious). Other parameters as in Fig. 2.

