## Supplementary material for "Adaptive ratchets and the evolution of molecular complexity": immune_dynamics.py

```
#######################################################################
# Survival of the fastest and the evolution of molecular complexity 
# Code for stochastic evolution of immune response
# Output presented in Figure 5 of the main text.
#######################################################################

import numpy as np
import pandas as pd
import matplotlib.pyplot as plt
import scipy as scipy
import pickle
import time
from tqdm import tqdm


def binding_affinity(x, a, b):
	
    return 1/(1+((np.exp(a+b*x))))

n0 = 2
e0 = 2.5
N_epitopes = np.arange(1, 10, 1)
rhos = np.logspace(np.log10(0.08), np.log10(2), 6)
tau = 1
T = 10000
R_bar = np.zeros((len(N_epitopes), len(rhos)))

for r, rho in enumerate(rhos):
	print('\n rho=%.1e \n'%(rho))
	for g in tqdm(N_epitopes):
	    n_mutations = np.zeros(g) #Array with number of mutatios in each epitope
	    neutralizations = binding_affinity(n_mutations, -n0*e0, e0) #Array with individuals neutralization probabilities
	    time = np.array([0]) #Array with time
	    t = time[-1]
	    time_intervals = np.array([])
	    exposure_times = np.array([0])
	    sick_times = np.array([])
	    R = np.array([1-np.product(1-neutralizations)]) #Array with total recognition function
	    propensities = np.array(np.concatenate(([rho]*g, [1/tau]))) #Array with propensities
	    i = 0
	    while (t<T): # Run Gillespie simulation until time T.
	        i+=1
	        cumsum = np.cumsum(propensities)
	        alpha = np.sum(propensities)
	        r1 = np.random.rand()
	        dti = (1/alpha)*np.log(float(1/r1))
	        time_intervals = np.append(time_intervals, dti)
	        t = t+dti
	        time = np.append(time, t)
	        r2 = np.random.rand()
	        idEvent = np.searchsorted(cumsum,r2*alpha)
	        #which event occured?
	        if(idEvent<g):#mutation
	            n_mutations[idEvent]+=1
	        else:#exposure
	            exposure_times = np.append(exposure_times, t)
	            r_sick = np.random.rand()
	            if(r_sick>R[-1]): #sick
	                n_mutations = np.zeros(g) #restart mutations
	                sick_times = np.append(sick_times, t)
	        neutralizations = binding_affinity(n_mutations, -n0*e0, e0) #update neutralizations  
	        R = np.append(R, 1-np.product(1-neutralizations)) #update R
	    if(g==1):
	    	R_bar_1 = 1-np.size(sick_times)/np.size(exposure_times)
	    #R_bar = np.append(R_bar, np.sum(R[:-1]*time_intervals)/time[-1])
	    R_bar[g-1, r] = (1-np.size(sick_times)/np.size(exposure_times))/1

df_R = pd.DataFrame(R_bar)
df_R.to_csv('./R_bar.txt', sep = '\t', index = False, header = False)
```
